# Protein Kinase A Negatively Regulates Ca^2+^ signaling in *Toxoplasma gondii*

**DOI:** 10.1101/265371

**Authors:** Alessandro D. Uboldi, Mary-Louise Wilde, Emi A. McRae, Rebecca J. Stewart, Laura F. Dagley, Luning Yang, Nicholas J Katris, Sanduni V. Hapuarachchi, Michael J Coffey, Adele M. Lehane, Cyrille Y Botte, Ross F. Waller, Andrew I. Webb, Christopher J. Tonkin

**Author notes:** Correspondence: Chris Tonkin, Division of Infection and Immunity, The Walter and Eliza Hall Institute of Medical Research, 1G Royal Pde, Parkville.

## Abstract

The phylum Apicomplexa comprises a group of obligate intracellular parasites that alternate between intracellular replicating forms and actively motile extracellular forms that move through tissue. Parasite cytosolic Ca^2+^ signalling activates motility, but how this is switched off after invasion is not understood. Here we show that the cAMP-dependent Protein Kinase A catalytic subunit 1 (PKAc1) of *Toxoplasma* is responsible for suppression of Ca^2+^ signalling upon host cell invasion. We demonstrate that that PKAc1 is sequestered to the parasite periphery by dual acylation of its regulatory subunit PKAr1. Newly invaded PKAc1-deficient parasites exit host cells shortly thereafter in a perforin-like protein 1 (PLP-1)-dependent fashion. We demonstrate that loss of PKAc1 results in an inability to rapidly downregulate cytosolic Ca^2+^ levels shortly after invasion. Furthermore, we demonstrate that PKAc1 also specifically negatively regulates resting cytosolic Ca^2+^ in conditions that mimic intracellularity. We also show that cAMP and cGMP have opposing role in microneme secretion, further supporting evidence that cAMP signalling has a suppressive role during motility. Together, this work provides a new paradigm in understanding how *Toxoplasma* and related apicomplexan parasites regulate infectivity.

## Introduction

The phylum Apicomplexa comprises a large group of obligate intracellular parasites that cause many important human and livestock diseases and includes *Plasmodium* spp. (malaria), *Cryptosporidium* spp. (severe diarrhoea) and *Toxoplasma gondii* (toxoplasmosis). *Toxoplasma* is transmitted to humans by eating undercooked meat harbouring cyst forms or by consuming soil, vegetables or water contaminated with oocysts shed from an infected cat. In healthy individuals, acute toxoplasmosis manifests with mild flu-like symptoms and is self-resolving, however, infection can cause life-threatening illness in the immunocompromised. Infection during pregnancy can cause abortion early in gestation or severe neurological developmental abnormalities in the foetus. Vertical transmission is also considered the cause of the relatively high rates of progressive blindness in some countries^1^.

*Toxoplasma*, like all apicomplexan parasites, critically requires the ability to move through tissue and invade host cells for survival and proliferation. Parasite movement is powered by a distinctive form of cellular locomotion referred to as gliding motility. Gliding motility is powered by the glideosome, an actomyosin-dependent motor located underneath the plasma membrane ^2^. Upon activation signals, single-pass transmembrane adhesins are released from the microneme organelles at the apical tip of the parasite and deposited onto the parasite surface. The current model suggests that motility is then driven by dynamic actin filaments attaching to the cytoplasmic tails of transmembrane adhesins via the Glideosome Associated Connector (GAC) ^3^, which are driven rearwards by the force generated by MyoA, thus resulting in forward directional movement of the parasite through tissue and into host cells ^2^.

Parasite egress, motility and host cell invasion is under tight control of intracellular signal transduction pathways. Both microneme and glideosome activity are controlled by cytosolic Ca^2+^ ([Ca^2+^]_cyt_) levels, cyclic GMP (cGMP) and inositol signalling pathways ^4–6^. The current model suggests that cGMP signalling is activated by unknown extracellular signals and results in stimulation of inositol phosphate metabolism and subsequent release of Ca^2+^ from intracellular stores as well as influx of Ca^2+^ from the external environment ^7,8^. A rise in [Ca^2+^]_cyt_ is temporally linked to the activation of egress from host cells and subsequent bursts of extracellular motility^9–11^. A rise in [Ca^2+^]_cyt_ then triggers the activation of Ca^2+^ -Dependent Protein Kinases (CDPKs) and a calcineurin phosphatase, which then phosphorylate/dephosphorylate substrates, triggering downstream events including activation of the glideosome and the release of adhesins from the micronemes ^7,12,13 14–19^.

To transition from immotile replicating tachyzoites to actively motile and invasive forms, parasites must sense the extracellular environment to regulate motility. High extracellular [K^+^], as encountered in the host cell, inactivates motility, whilst a drop in this ion causes a rise in [Ca^2+^]_cyt_ and activation of locomotion ^20^. More recently, it has been shown that extracellular pH ([H^+^]) also regulates motility. Here, low extracellular pH potently activates motility, whilst also being able to overcome the suppressive effect of high extracellular [K^+^] ^21^. During intracellular growth, a lowering of the pH of the vacuolar space activates Ca^2+^-dependent egress, whilst also promoting an acidic environment for the activation of Perforin-Like Protein 1 (PLP-1) to elicit membrane damage and tachyzoite egress ^21^.

Despite advances in our understanding of how *Toxoplasma* and other apicomplexan parasites activate motility, it remains unclear how these important pathogens sense environmental cues. In other eukaryotic systems G-Protein Coupled Receptors (GPCRs) are essential for environmental sensing and act by receiving extracellular cues and transducing these signals across the plasma membrane. GPCRs physically associate with signalling proteins on the cytoplasmic side of the plasma membrane, causing changes in their activity upon extracellular cues. GPCRs commonly activate cyclic adenosine monophosphate (cAMP) signalling pathways. This occurs by direct coupling of GPCRs to adenyl-cyclases, which are activated to produce cAMP, upon encountering environmental stimuli. cAMP then activates a tetrameric Protein Kinase A (PKA) complex, by binding to the regulatory subunit (PKAr), promoting its dissociation from the catalytic subunit (PKAc), thus promoting downstream phosphorylation. Previous work has suggested that PKA, and thus cAMP signalling, operates in Apicomplexa and is important for parasite invasion in *Plasmodium*, in part through the phosphorylation of the cytoplasmic tail of the parasite adhesin AMA1^22,23^ and may be linked to Ca^2+^ signalling^13^. Interestingly, one of three PKA paralogues in *Toxoplasma* negatively regulates bradyzoite differentiation ^24^.

Here we investigate the function of PKAc1 in *Toxoplasma* and show that this kinase acts as a negative regulator of tachyzoite egress and is required for the transition of invasive tachyzoites to form intracellular replicative forms. Whilst PKAc1-depleted tachyzoites are able to invade normally and form a parasitophorous vacuole (PV), they rapidly egress host cells shortly thereafter. We show that in the absence of PKAc1, newly invaded tachyzoites cannot rapidly quench [Ca^2+^]_cyt_ and this leads to an overall higher resting level. Furthermore, we provide evidence that cAMP and cGMP have opposing role in regulating microneme secretion, where cAMP dampens exocytosis of these organelles. Together, our work pinpoints a critical role of PKAc1, and thus cAMP signalling, in negatively regulating motility by inhibiting cytosolic Ca^2+^ levels upon parasite invasion. Our work therefore, provides the evidence of how *Toxoplasma*, and potentially other apicomplexan parasites, switch-off motility to begin intracellular replication.

## Results

### A PKA complex in *Toxoplasma* localises to the parasite periphery

We wished to understand if cAMP/PKA signalling participates in relaying environmental cues, to modulate motility. *Toxoplasma* contains three annotated PKA catalytic subunit paralogues (toxodb.org) ^25^. Previous work has shown that PKAc3 (TGME49_286470) negatively regulates the differentiation of bradyzoite forms during tachyzoite growth ^24^, whilst PKAc2 (TGME49_228420) appears not expressed in tachyzoite stages (toxodb.org). PKAc1 (TGME49_226030) is the most widely conserved *Toxoplasma* paralogue across the Apicomplexa and thus was chosen for study. To understand the function of PKAc1 we genetically introduced an epitope tag at the 5’ end of the protein, whilst simultaneously putting the gene under conditional genetic control using the tet-off system (Figure S1) (see below for description). To assess the localisation of PKAc1 we performed an immunofluorescence assay (IFA) using anti-HA antibodies and observed a peripheral staining pattern, as marked by IMC1 antibodies (Figure 1Ai). In some tachyzoites we noticed internal staining, which co-localised with the apical inner membrane complex (IMC) marker ISP1, suggesting association with the forming daughter cells during replication (Figure 1Aii).

**Figure 1:**
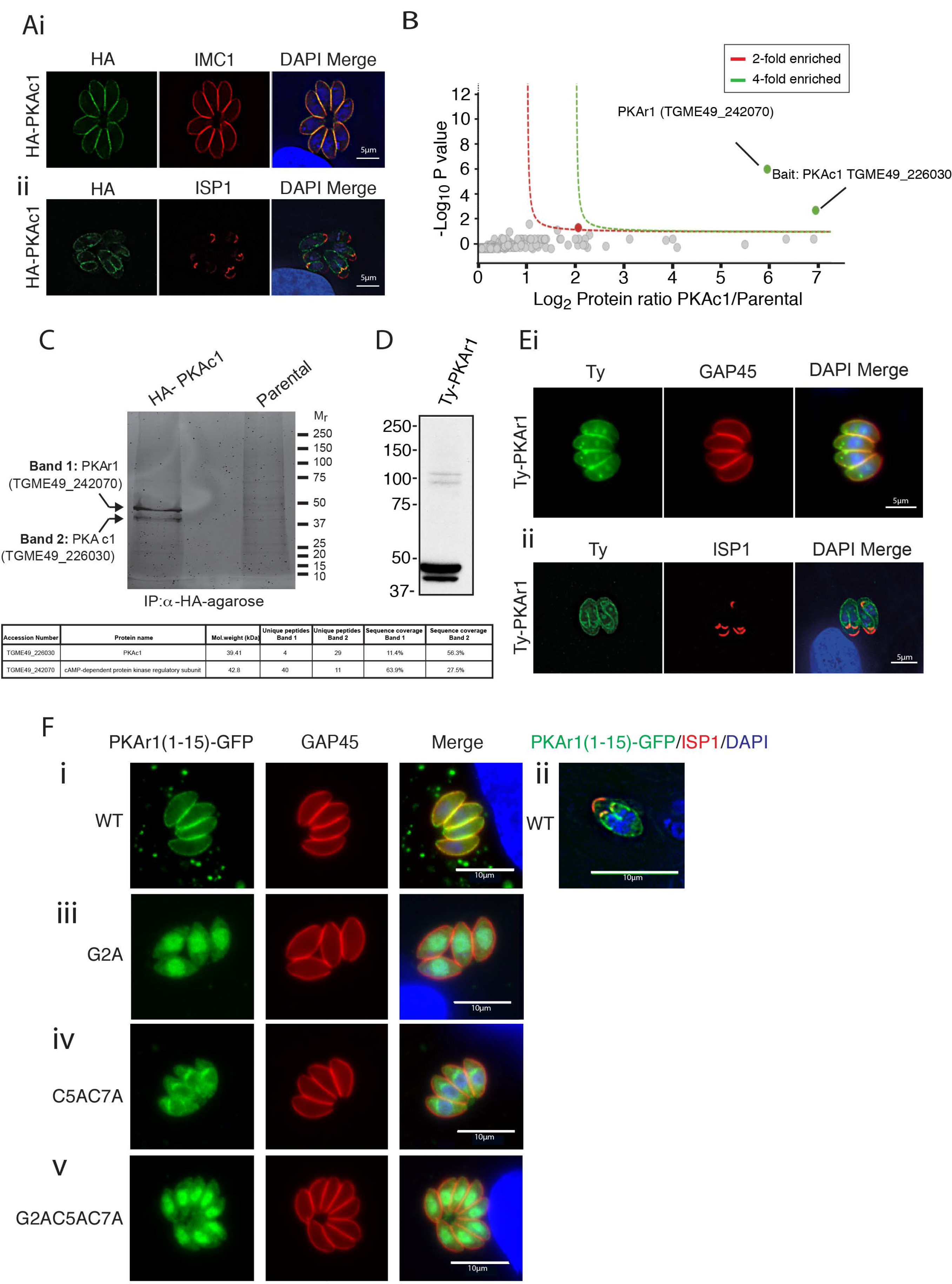
*Toxoplasma* PKAc1 localises to the tachyzoite periphery likely by dual acylation of its regulatory subunit PKAr1. (A) Localisation of HA-tagged PKAc1 by IFA, co-staining with the peripheral marker IMC1 (i) and the apical IMC marker -ISPl(ii). (B) Identification of PKAc1 interacting proteins by anti-HA immunoprecipitation whole eluate tryptic digestion, mass spectrometry and quantitation using spectral counting. Red and green lines signify 2-fold and 4-fold enrichment, as compared to the parental line (see Table S2 for full list of identified proteins). (C) In gel digest and mass spectrometry, where table denotes proteins and coverage identified in each band. (D) Western blot of Ty epitope-tagged PKArl (TGME49_242070). (E) Localisation of Ty epitope tagged PKArl by IFA, co-stained with either (i) the peripheral marker GAP45 or (ii) the apical IMC protein ISPl, outlining localisation to the developing daughter cells. (F) Mutational analysis of the role of putative myristoylation and palmitoylation sites at the N-terminus of PKArl in membrane sequestration. (i) The first l5 amino acids of PKArl (PKArl(l-l5)_WT_) were fused to GFP and co-localised with GAP45 and (ii) ISTl, showing localisation to the periphery and to the developing daughter cells, respectively. (iii) Putative myristoylation site glycine at position 2 was mutated to alanine (PKArl(l-l5)_G2A_), (iv) Putative palmitoylated cysteine residues at position 5 and 7 were mutated to alanine ‒(PKArl(l-l5)_C5AC7A_). (v) Both putative myristoylation and palmitoylation sites were mutated ‒(PKArl(l-l5)_G2AC5AC7A_). In each case localisation of GFP ‒fusions were compared to the peripheral marker GAP45 and counterstained with DAPI.

Across eukaryotes PKA typically localises to defined cellular compartments via association with A-kinase Associated Proteins (AKAP) and/or by binding to GPCRs. To determine if this was true in *Toxoplasma* we performed immunoprecipitation of HA-tagged PKAc1 and compared eluates to parental controls either by in-solution by trypsin digestion (Figure B) or in gel digests, followed by mass spectrometry to detect associated proteins (Figure 1C). In both cases we only confidently detected one additional protein, which we determined to be TGME49_242070, a protein with several predicted cAMP-binding domains, and thus likely a PKA regulatory subunit (henceforth referred to as PKAr1).

PKAr subunits in other eukaryotes contain an N-terminal docking domain that is responsible for associating the PKA complex with AKAP. *Toxoplasma* PKAr1 appears to lack this domain and instead contains a glycine at position 2 and two cysteines at positions 5 and 7, typical of sites of acylation. Acylation is an important mechanism for imparting membrane affinity on a range of proteins involved in signalling and motility in *Toxoplasma* ^26–28^. To investigate the localisation of PKAr1 without disturbing this potential motif we introduced a Ty epitope tag 15 amino acids downstream from the starting methionine. Western blot of Ty-PKAr1 revealed that there were two major species of this protein close to the expected size, the upper species being more abundant and of a size consistent with a modified (e.g. acylated) form (Figure D). There is also two bands at ∼100kD, but the identity or significance of these is not known. Co-localisation of Ty-PKAr1 by IFA with GAP45 also demonstrated a largely peripheral localisation and further, we also observed localisation to the nascent IMC (as marked by ISP1) during internal daughter cell budding (Figure Ei and ii), similar to that observed with PKAc1. PKAr1 also has apparent internal pools separate from the IMC, one of which localises within the nucleus (Figure Ei), the significance of this is unknown.

PKAr1 was recently described to be part of the *Toxoplasma* ‘palmitome’, thus further supporting the evidence that this protein is acylated ^29^. To investigate whether the N-terminal region of PKAr1 was responsible for sequestration to the periphery by myristoylation of glycine 2 and palmitoylation of cysteine 5 and/or 7, we fused the first 15 amino acids of PKAr1 to GFP and used IFA to monitor localisation. We found that, indeed, PKAr1(1-15) was sufficient to target GFP to the parasite periphery (Figure 1Fi) and to internal budding daughter cells (Figure 1Fii). Furthermore, mutation of the putative myristoylation site to an alanine (G2A) resulted in a severe abrogation of peripheral localisation and instead caused an accumulation of the GFP fusion protein in the cytoplasm and nucleus (Figure 1Fiii). Mutation of the two putative palmitoylation acceptor cysteines to alanines (C5A C7A), also resulted in abrogation of peripheral localisation (Figure 1Fiv) as did mutation of all three residues simultaneously (G2A C5A C7A) (Figure 1Fv). Together, these data are consistent with PKAc1 localising to the parasite periphery, likely the IMC membranes, via its association with PKAr1, which relies on dual acylation to derive membrane avidity.

### PKAc1 is essential for the tachyzoite lytic cycle

To investigate the function of PKAc1 in *Toxoplasma* tachyzoites we generated a PKAc1 conditional knockout (cKo) line by replacing the endogenous promoter with the tet-off (T7S4) promoter (Figure S1) (Figure 2A) ^30,31^. To monitor the regulation of PKAc1 in the PKAc1 cKO parasites we tracked protein production using HA antibodies. Treatment of parasite cultures with anhydrotetracycline (ATc) for 48h led to a marked decrease in PKAc1 levels (Figure 2B), and IFA showed loss of detectable levels of this protein (Figure 2C). To monitor growth of PKAc1 cKO parasites across the complete lytic cycle we then performed plaque assays on confluent monolayers of HFF. ATc treatment of PKAc1 cKO, but not parental, parasites resulted in a drastic reduction in plaque size, consistent with this kinase having an important role in one or more steps of the *Toxoplasma* lytic cycle and furthermore is congruent with its deleterious CRISPR screen growth score ^32^ (Figure 2D). During our preliminary analysis, we noticed that in vitro cultures of PKAc1-depleted tachyzoites growing in host cells had an unusual presentation. After 24 h of ATc treatment, PKAc1 cKO cultures contained very few intracellular parasites (compared to untreated cultures). Concurrently, host cells appeared to have been lysed, some detaching form the surface plastic (Figure 2E and S2).

**Figure 2:**
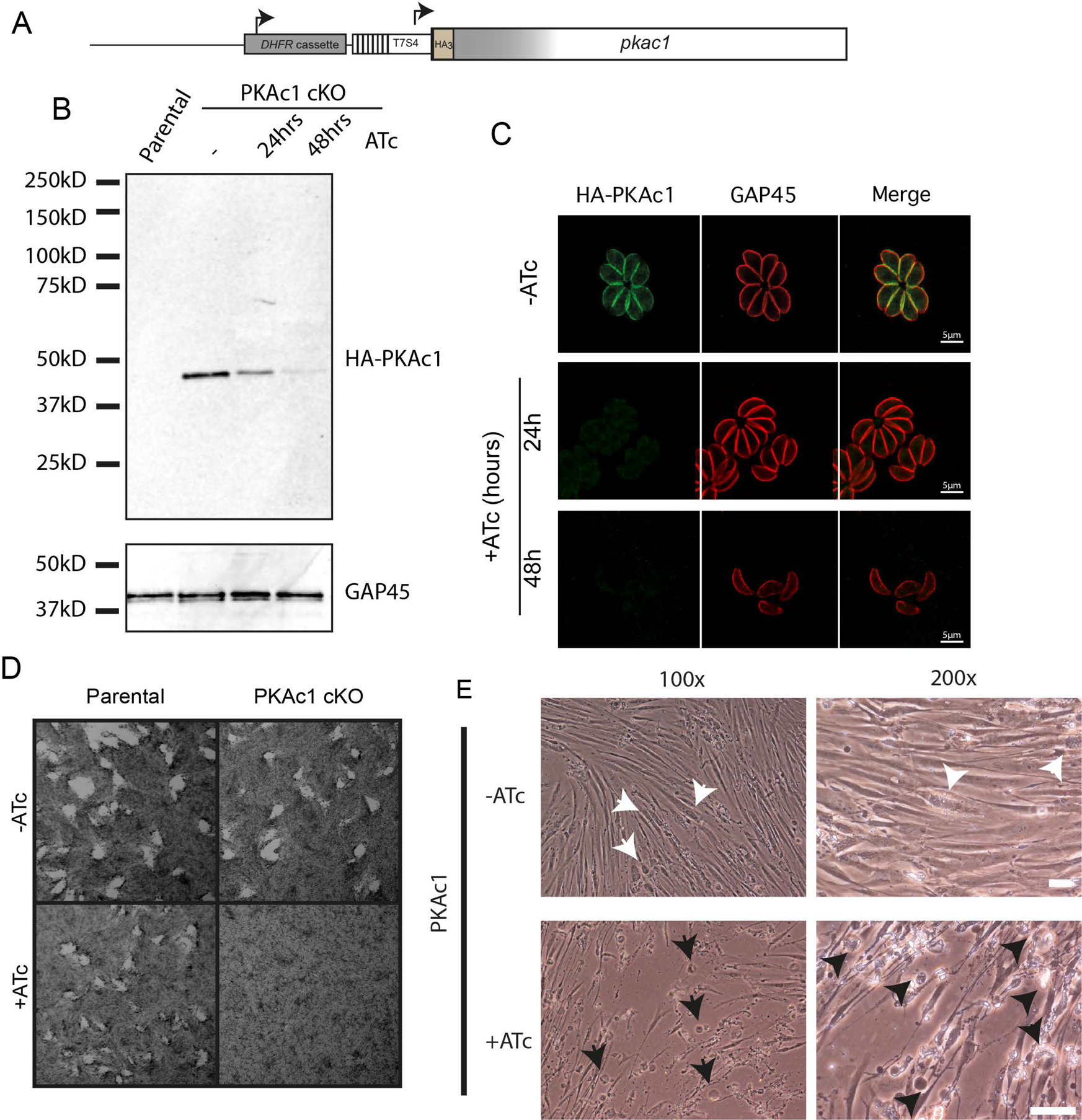
Generation and characterization of a conditional knockout of *Toxoplasma* PKAc1. (A) Schematic representation of tet-off promoter replacement-based conditional knockout of PKAc1 (PKAc1 cKO). Details of genetic strategy and validation of PKAc1 cKO are available in Figure S1. (B). Downregulation of PKAc1 levels upon ATc treatment at 24hrs and 48rs and comparison to parental line. HA antibodies were used to detect PKAc1 and GAP45 antibodies were used as a loading control. (C) IFA of PKAc1 upon ATc treatment for 24hrs and 48hrs. (D) Plaque assay of parental and PKAc1 cKO and parental lines +/− ATc upon infection of confluent HFF monolayers. (E) Images of in vitro cultures of HFF infected with PKAc1 cKO tachyzoites after for 24 hours growth +/− ATc. White arrows point to intact intracellular parasite vacuoles and black arrow show examples of unhealthy and detached HFFs. Scale bars = 50μm. More images are available in Figure S2.

### PKAc1 is required for stable invasion of host cells

We hypothesised that the unusual presentation of PKAc1-deficient- host cell cultures was due to aberrant invasion. We therefore performed two-colour invasion assay to enumerate invasion efficiency on a population level. After a 10-minute invasion period we observed a severe loss of intracellular parasites (Figure 3A). To get a better understanding of the role of PKAc1 during invasion we undertook live cell imaging. In ATc-treated or untreated parental parasites and untreated PKAc1 cKO tachyzoites we saw typical invasion, where parasites attached to host cell monolayers, activated motility and penetrated the host cell through a tight constriction, typical of the formation of the moving junction (Figure 3Ai, ii, iii, Movie S1, S2, S3). Following invasion, tachyzoites remained stationary and little movement of the host cell was observed (invading tachyzoites marked with white arrow head and dashed white lines mark outline of host cell) (Figure 3Ai, ii, iii, Movie S1, S2, S3). Upon depletion of PKAc1, shortly after the apparent completion of invasion, we saw that the host cell began to detach from glass slide surface often with the concomitant re-activation of parasite motility (Figure 3Ci, ii, Movie S4, S5). We observed that tachyzoites could directly exit host cells (Figure 3Cii, Movie S4), and in some cases, move within the confines of the damaged host cell (Figure 3Ci)(Movie S5). We then quantified the timing of host cell collapse in relation to invasion over a larger population of cells. Over a period of 10 minutes of filming we saw no parental (+ATc) or PKAc1 cKO (–ATc) egress (Figure 3D). In comparison, some PKAc1-deficient tachyzoites exited as quickly as ∼30 seconds post invasion, whereas others did not exit within the 10- minute filming period. Over the population, host cell collapse occurred an average of ∼200 seconds post invasion (Figure 3C). We then quantified host cell damage on a population level, as a function of parasite concentration by monitoring host cell integrity using crystal violet staining. To do this, ATc treated and untreated parental and PKAc1 cKO tachyzoites were serially diluted and allowed to invade host cells for 3h, after which time host cell integrity was analysed by crystal violet staining (Figure 3E). PKAc1-depleted tachyzoites caused a decrease in the number of intact host cells, with the damage to the host cell monolayer more prominent with increasing parasite number. In contrast, PKAc1 cKO parasites expressing PKAc1 and parental parasites treated with ATc caused no such damage (Figure 3E). Together, these results suggest that PKAc1 is required for productive invasion and/or the suppression of motility once cell entry is complete.

**Figure 3:**
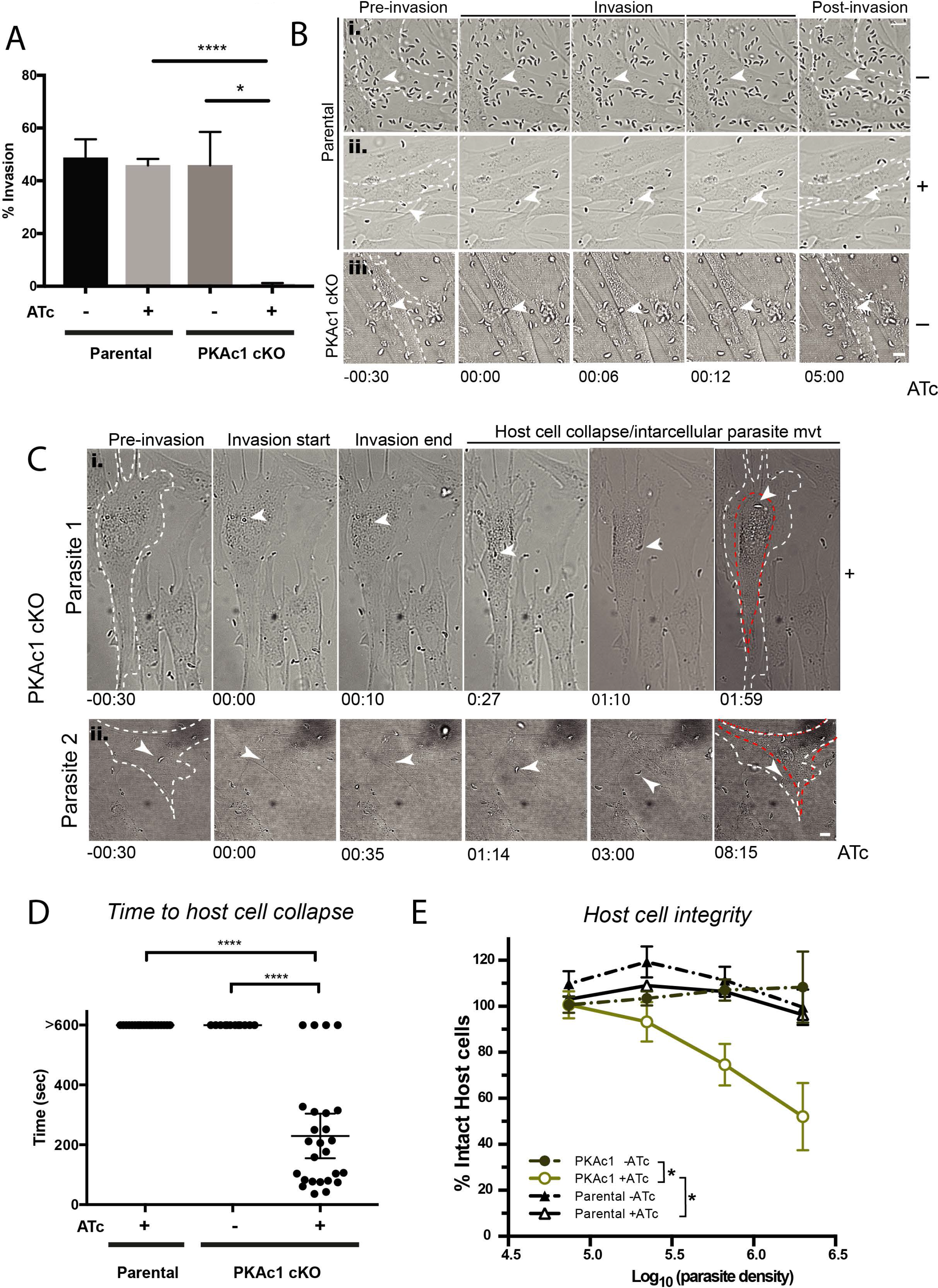
PKAc1-deficient *Toxoplasma* tachyzoites damage host cells upon invasion. (A)Static invasion assay of parental and PKAc1cKO tachyzoites. N=3 and Data is the mean ± SEM of 3 independent experiments. For each replicate, parasites in 16 panels, each with an area of 211.3 × 198.1 microns were counted at 63 X magnification. P values are calculated using an unpaired two-tailed t-test, where *= <0.05 and **** =<0.0001. (Bi and ii) Live cell imaging of parental strain +/− ATc and, (iii) PKAc1 cKO without ATc treatment, before, during and after invasion of host HFFs. White arrow heads point to invading tachyzoites and dashed white lines signify the border of the target HFF before and after invasion. Time in minutes and seconds is displayed across the bottom of panels. Stills correspond to Movies S1, S2 and S3, respectively. Scale bar = 10μm, (Ci and ii) Live cell imaging of two separate PKAc1-depleted tachyzoites (+ATc), before, during and after invasion of HFF. As above, white dashed lines outline host cell border pre-invasion, whilst red border denotes HFF outline post invasion. White arrows signify tracked tachyzoites. Time in minutes and seconds is displayed across the bottom of panels. Stills correspond to Movie S4 and S5, Scale bar = 10μm (D) Quantification of time to host cell collapse in Parental and PKAc1-depleted lines. >600 signifies that host cell collapse was not observed over the 10-minute (600 second) filming period. Data represents mean ± 95% confidence intervals, Parental + ATc n= 20, PKAc1-ATc n=12, PKAc1 +ATc n=26, across at least 3 experiments. P values are calculated using an unpaired two-tailed t-test, where **** =<0.0001. (E) Integrity of host cells invaded for 1 hour by parental and PKAc1cKO lines, +/− ATc as measured as a function of tachyzoite number versus crystal violet absorbance. Data represents mean ±SEM over 4 independent experiments. P values are calculated using a two-way ANOVA, where *=<0.05 only at highest tachyzoite concentration.

### PKAc1 does not play a detectable role in invasion

The exit of host cells by PKAc1 depleted tachyzoites shortly after invasion could be caused either by a defect in the formation or sealing of the parasitophorous vacuole membrane (PVM) during invasion, or by parasites activating egress shortly after invasion is complete. To test if PKAc1 has a role in invasion, we measured the speed of invasion (Figure S3A) and the ability of invading tachyzoites to form a moving junction (as marked by RON4) (Figure S3B). In both cases we saw no difference between parental and PKAc1-depleted lines (representative images shown of RON4 staining). We then determined whether PKAc1 plays a role in PVM formation. Secretion of rhoptries is required for PVM formation and can be measured by inhibiting invasion using cytochalasin D (CytD) and staining for the presence of ‘empty vacuoles’ (evacuoles) secreted into the host cell ^33^. We quantitated evacuole production using ROP1 as a marker. Further, we used MIC8 cKO as a positive control as this protein is known to be important for their production ^34^. Here we observed that PKAc1-depleted tachyzoites were able to produce evacuoles at the same rate as parental lines, suggesting that this kinase plays no role in the secretion of rhoptry contents (Figure S3C). We also measured formation of the PVM on invading and invaded parasites using a mouse embryotic fibroblast (MEF) line that expresses membrane-bound tdtomato ^35^. In static images, where PKAc1 and parental lines were expressing cytosolic GFP we observed clear formation of an intense tdtomato+ around the body of tachyzoites, as clearly demonstrated by plotting intensity values across a cross section (Figure S3Di and ii). Furthermore, when performing live cell imaging on tachyzoites invading tdtomato-expressing MEFs we saw formation of a tdtomato^+^ membrane surrounding invading parasites, which persisted until collapse of the host cell (Figure S3E and Movie S6 for representative time-lapse), further suggesting the formation of a PVM even in the absence of PKAc1.

We also assayed whether PKAc1 was involved in motility, host cell attachment as well as microneme secretion (Figure S4). To assay tachyzoite motility we performed live cell imaging and quantitated total motile and non-motile fractions as well segregating based on the different types of motility when put in intracellular (IC) or extracellular (EC) buffer. Whilst we found an increase in the fraction of motile tachyzoites in EC buffer we saw no changes upon repression of PKAc1 expression (Figure S4Ai and ii). We also measured tachyzoite host cell attachment using standard assays, and again, found no differences of PKAc1-deficient parasites as compared to controls (Figure S4B). Furthermore, we assayed secretion of micronemal proteins into the supernatant using both quantitative western blot and quantitative proteomics. In both cases we found that PKAc1-depleted parasites had no difference to parental and no ATc controls (Figure S4C, D and E). Note that when performing quantitative proteomics on supernatants we did see more peptides from proteins known or predicted to be found inside the parasite. Whilst we do not know the reasons behind this finding we suggest that loss of PKAc1 may make the plasmamembrane more fragile and prone to lysis (Figure S4E). Together these results suggest that PKAc1 has no detectable role in invasion, motility or host cell adhesion.

### Perforin-like Protein 1 is required for premature host cell egress in PKAc1 deficient tachyzoites

We then wanted to determine whether we could determine whether the PKAc1-deficient phenotype was a reliant on programmed host cell egress. *Toxoplasma* PLP-1 is critical to induce PVM and host cell membrane breakdown required for parasite egress ^36,37^ Furthermore, vacuolar acidification is required for activating PLP-1 ^21^ to elicit membrane damage. We therefore wondered whether early egress of PKAc1-deficient tachyzoites requires acidification of the vacuolar space and the cytolytic activity of PLP-1. Furthermore, given that PLP-1 has no role in invasion, testing for dependency on PLP-1 and vacuolar acidification of the premature egress phenotype of PKAc1-depleted parasites would support a role for this kinase as a negative regulator of egress post-invasion. To test this, we first genetically deleted PLP-1 in our PKAc1 cKO line (Figure S5). Western blot of PKAc1 cKO/Δ*plp-1* parasite lysates using PLP-1 antibodies showed a loss of signal verifying gene disruption (Figure 4A). To determine if PLP-1 is required for early egress of PKAc1-deficient parasites we grew PKAc1 cKO and PKAc1 cKO/Δ*plp-1* tachyzoites in the presence of ATc for 24 h and first monitored the gross morphology of parasite infected host cells under the microscope. As previously shown in Figure 2, loss of PKAc1 results in unhealthy-looking host cells that appear to readily detach from the substrate with no intracellular parasites evident (Figure 4B-black arrows). Conversely, host cells infected with parasites also lacking PLP-1 (PKAc1 cKO/Δplp-1 +ATc) appeared to have a reversal of this effect, containing many late-stage vacuoles full of tachyzoites, similar to parental and non-ATc treated lines (Figure 4B- white arrows and Figure S2). This suggests that loss of this cytolytic protein at least partially rescues genetic depletion of PKAc1. To quantify the role of PLP-1 in host cell destruction post invasion in PKAc1-deficient tachyzoites we performed an end-point invasion assay. As compared to the loss of PKAc1 alone, (as reproduced from above), the additional deletion of PLP1 resulted in the presence of significantly more intracellular parasites at 10 minutes post invasion (Figure 4C).

**Figure 4:**
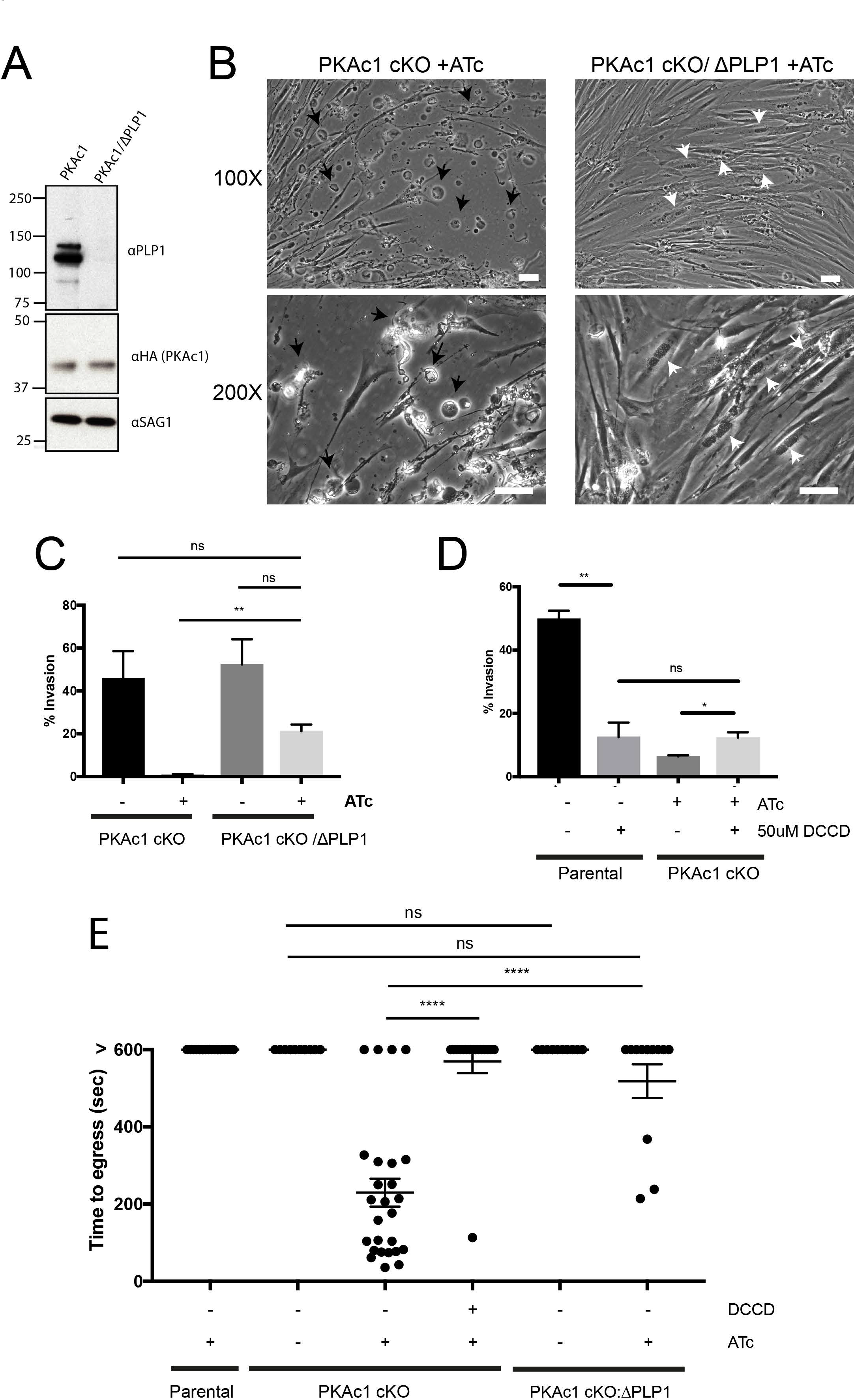
PLP-1 controls host cell damage in PKAc1-deficient tachyzoites. (A) Genetic deletion of PLP-1 (ΔPLP1) in PKAc1 cKO as confirmed by western blot analysis. See Figure S5 for genetic strategy and genotyping (B). Morphology of PKAc1- and PKAc1/PLP-1- deficient tachyzoites during in vitro growth in HFF after 24 hrs of ATc treatment. White arrows point to intracellular tachyzoites after several rounds of replication. Black arrow point to examples of collapsed host cells. More images in Figure S2. (C) Static invasion assay (after 10 minutes) of PKAc1 cKO/ΔPLP1 tachyzoites as compared to PKAc1 cKO (reproduced from Figure 3). (D) Effect of treatment with DCCD on host cell invasion of parental and PKAc1 cKO ±ATc. Data from C and D represents mean ± SEM from 3 independent experiments. P values are calculated using an unpaired two-tailed t-test, where *=<0.05, ** =<0.01, and ns= not significant. (E) Quantification of time to host cell collapse in Parental and PKAc1and PKAc1/ΔPLP1tachyzoites ±ATc and ± 50μM DCCD. >600 signifies that host cell collapse was not observed over the 10-minute (600 second) filming period. Parental +ATc, PKAc1 ±ATc data is reproduced from figure 3 for purposes of comparison. Data represents mean ± 95% confidence intervals over at least 3 independent experiments where PKAc1 +ATc +50μM DCCD n=16, PKAc1/ΔPLP1 -ATc n= 7, PKAc1/APLP1 +ATc n= 12.

Previously, it has been shown that the inhibitor of H^+^ ATPase transporters N,N’- Dicyclohexylcarbodiimide (DCCD) can, prevent vacuolar acidification and inhibit PLP-1 activity. We therefore also tested whether treatment of DCCD could reverse the loss of PKAc1, thus further supporting a role for this kinase in supressing egress post invasion. First, using a standard invasion assay we could show that whilst DCCD significantly inhibits invasion of PKAc1-expressing parasites, it also however, rescues the proportion of observable intracellular parasites of PKAc1-depleted tachyzoites after 10 minutes of invasion, suggesting a reversal of the host cell damaging effects upon loss of this kinase. Further, there was no difference in amount of intracellular tachyzoites with and without PKAc1 that were both treated with DCCD (Figure 4D).

To confirm these findings and further dissect the PKAc1-depletion phenotype dependency on PLP-1 and acidification we performed live cell imaging on PKAc1 cKO/Δplp-1 tachyzoites and quantitated how long tachyzoites remained intracellular following invasion. As compared to PKAc1-deficient parasites, the additional loss of PLP1 saw the vast majority of parasites remain intracellular over the 10 minutes of filming (Figure 4E). Tracking individual PKAc1-depleted tachyzoites treated with 50uM of DCCD also saw almost all parasites remain intracellular after invasion over the 10-minute filming period. Together, this data further supports a role of PKAc1 in negatively regulating host cell egress and demonstrates that PKAc1-dependent egress host cells occurs in a PLP-1-dependent fashion.

### PKAc1 controls rapid dampening of cytosolic Ca^2+^ after invasion

Tachyzoite [Ca^2+^]_cyt_ has been temporally and functionally linked to parasite egress, motility and invasion ^9,10 11,38^. We hypothesised that PKAc1 may function to negatively regulate [Ca^2+^]_cyt_, where loss of this kinase sustains parasite motility and enables host cell egress to occur. Previously, we have used the genetically-encoded biosensor – GCaMP6 – to monitor cytosolic levels of Ca^2+^ in intracellular, egressing and extracellular motile tachyzoites ^9,39^ To first assess the utility of GCaMP6 in monitoring Ca^2+^ levels during invasion we stably introduced the GCaMP6/mCherry expressing plasmid at the *uprt* locus into the parental line (Dku80:TATi) and quantitated fluorescence levels (+ATc treatment) over a 10 min period, acquiring 5 z-projections every ∼1.3 s (Figure 5, Movie S7). We observed that invasion of the parental line coincides with a rapid quenching of GCaMP6 fluorescence, dropping to 20-40% of maximum levels in a period of 10-20 s post invasion (t=0 = beginning of invasion, blue arrow=completion of invasion, red arrow = time to reach 35% of maximum fluorescence) (Figure 5Ai and ii, Movie S7). We found that this pattern was seen in all observable invasion events in the parental line (Figure 5Aiii, individual traces see Figure S6). We then introduced the GCaMP6/mCherry expressing plasmid into PKAc1 cKO parasites and measured the dynamics in fluorescence as a readout for changes in [Ca^2+^]_cyt_. In the absence of ATc, PKAc1 cKO tachyzoites were able to quench [Ca^2+^]_cyt_ shortly after invasion, similar to the parental line (Figure 5Bi,ii,iii, Movie S8, individual traces Figure S7). However, depletion of PKAc1 by ATc treatment resulted in drastic changes in Ca^2+^ dynamics. Here we noticed that PKAc1- depleted tachyzoites could not rapidly dampen GCaMP6 fluorescence after invasion was complete (green arrow = time of host cell collapse)(Figure 5Ci, ii). Instead, as viewed by individual traces of PKAc1 depleted tachyzoites, [Ca^2+^]_cyt_ varied greatly during this time (Figure 5Ciii and Figure S8). In order to quantitate and graphically represent cytosolic Ca^2+^ dynamics over time we chose, based on parental controls, a value of 35% of maximum fluorescence to baseline after the completion of invasion (Figure 5Aii, iii and Bii, iii). By measuring the time taken for tachyzoites to reach 35% of maximum we could observe that PKAc1-depleted parasites took significantly longer than untreated or parental controls to reduce [Ca^2+^]_cyt_, averaging ∼150 seconds to reach 35% as compared to controls which typically took 20-40 seconds (Figure 5D). We then measured GCaMP6 fluorescence level at 100 seconds, a time when all tachyzoites of control samples have completed invasion and dampened [Ca^2+^]_cyt_ (Figure 5A and B). Tachyzoites lacking PKAc1 expression were then split into those that egressed before and after t=100 as well as those that did not egress at all. In doing so we could see that those PKAc1-deficient parasites that did egress had significantly higher GCaMP6 fluorescence at this time point, whilst those that did not had levels equivalent to controls (Figure 5E). Overall, this work suggests that PKAc1 plays an important role in the rapid reduction of [Ca^2+^]_cyt_ that normally takes place shortly after invasion is complete.

**Figure 5:**
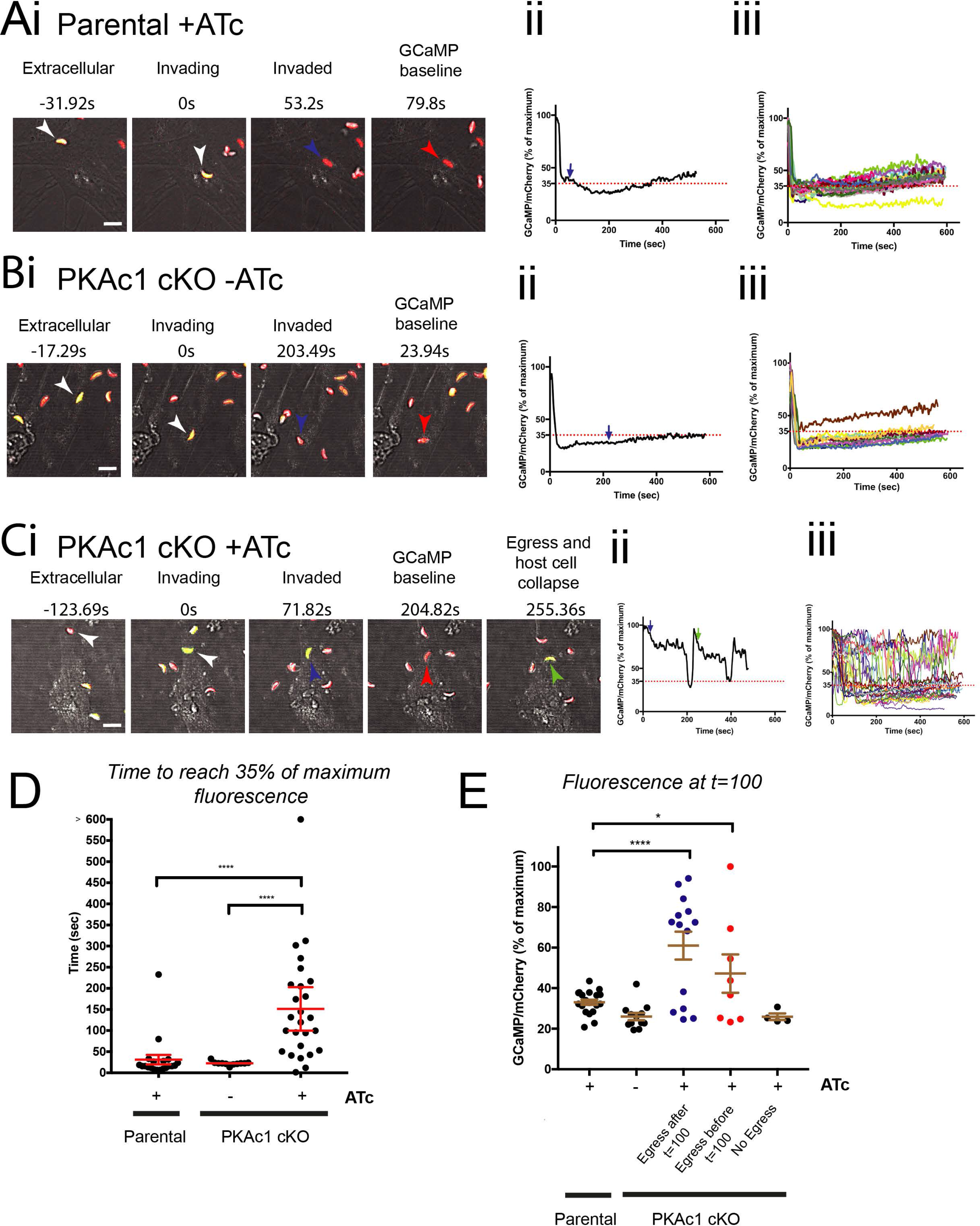
PKAc1 controls the rapid down-regulation of cytosolic Ca^2+^ shortly after invasion. (A) Live cell imaging of Parental cKO tachyzoites with ATc treatment, (B) PKAc1 without ATc treatment and (C) PKAc1 treated with ATc all stably expressing GCaMP6f/mCherry at the *uprt* locus. In A, B and C; (i) outlines images of individual movies, showing when parasites are extracellular, invading, upon completion of invasion (blue arrow) at ‘baseline’ (defined as when GCaMP/mCherry ratio, reaches 35%) (see accompanying Movies S7, S8 and S9, respectively). (ii) In each case, shows a graphical representation of tracked parasites in (i), where blue line denotes moment of completion of invasion and green arrow shows moment of host cell egress. Dotted lines marks 35% of maximum GCaMP/mCherry ratio, defined as ‘baseline’. (iii) In each case, shows overlays of all traced parasites. Figure S6, S7 and S8 show individual traces of each tracked parasite, respectively. (D) Graphical representation of each tracked parasite across all conditions showing time to ‘baseline’ fluorescence (as defined as 35% of maximu). (E) Graphical representation of normalised GCaMP/mCherry ratio at t= 100 seconds across all parasites in all conditions, and in the case of PKAc1 cKO +ATc, segregated depending on time of egress. Data in D and E is represented as mean ± 95% confidence interval. P values are calculated pairwise, using an unpaired two-tailed t-test, where *=<0.05, **** =<0.0001.

### Live cell imaging reveals that PKAc1 regulates cytosolic Ca^2+^ independent of PLP-1 activity

We wondered whether [Ca^2+^]_cyt_ in PKAc1-deficient parasites was influenced by PLP-1-dependent egress and exposure to extracellular environment. To test this, we stably introduced GCaMP6/mCherry expressing plasmid into the *uprt* locus of PKAc1 cKO/Δ*plp1* tachyzoites (Figure 6). In the absence of ATc, PKAc1 cKD/Δ*plp1* tachyzoites invaded host cells in a typical fashion, which was then followed by rapid dampening of [Ca^2+^]_cyt_, indistinguishable to parental controls (Figure 6Ai,ii), which was highly consistent across the population (Figure 6Aiii). Upon depletion of PKAc1 with ATc we saw that GCaMP6 levels did not dampen in a typical fashion, demonstrating that the cytolytic activity of PLP-1 and thus early exposure to the extracellular environment plays little role in the loss of cytosolic Ca^2+^ dampening in PKAc1- deficient parasites (Figure 6Bi, ii, iii). To further explore the independence of [Ca^2+^]_cyt_ from membrane disruption by PLP-1 we treated parasites with the H^+^ ATPase inhibitor DCCD (Figure 6C). After tracking many invading tachyzoites, we saw that DCCD treatment of PKAc1-deficient parasites largely phenocopied PLP-1 deletion in terms of loss of egress but sustained high [Ca^2+^]_cyt_ (Figure 6Ci,ii, iii), thus further supporting the notion that regulation of [Ca^2+^]_cyt_ is not influenced by cytolytic activity of PLP-1. We also quantitated these effects by graphing time taken of each treatment to reach 35% of maximum fluorescence (Figure 6D) as well as fluorescence levels at t=100 seconds (Figure 6E). As compared to PKAc1-depletion alone (as re-displayed here from Figure 5 here) this showed that PLP-1-deficient and DCCD treated PKAc1-deficient tachyzoites maintain a higher [Ca^2+^]_cyt_ level post invasion (Figure 6D and E). Together these data provide strong evidence that PKAc1 negatively regulates [Ca^2+^]_cyt_ post-invasion independent of PLP-1-dependent PVM and host cell lysis.

**Figure 6:**
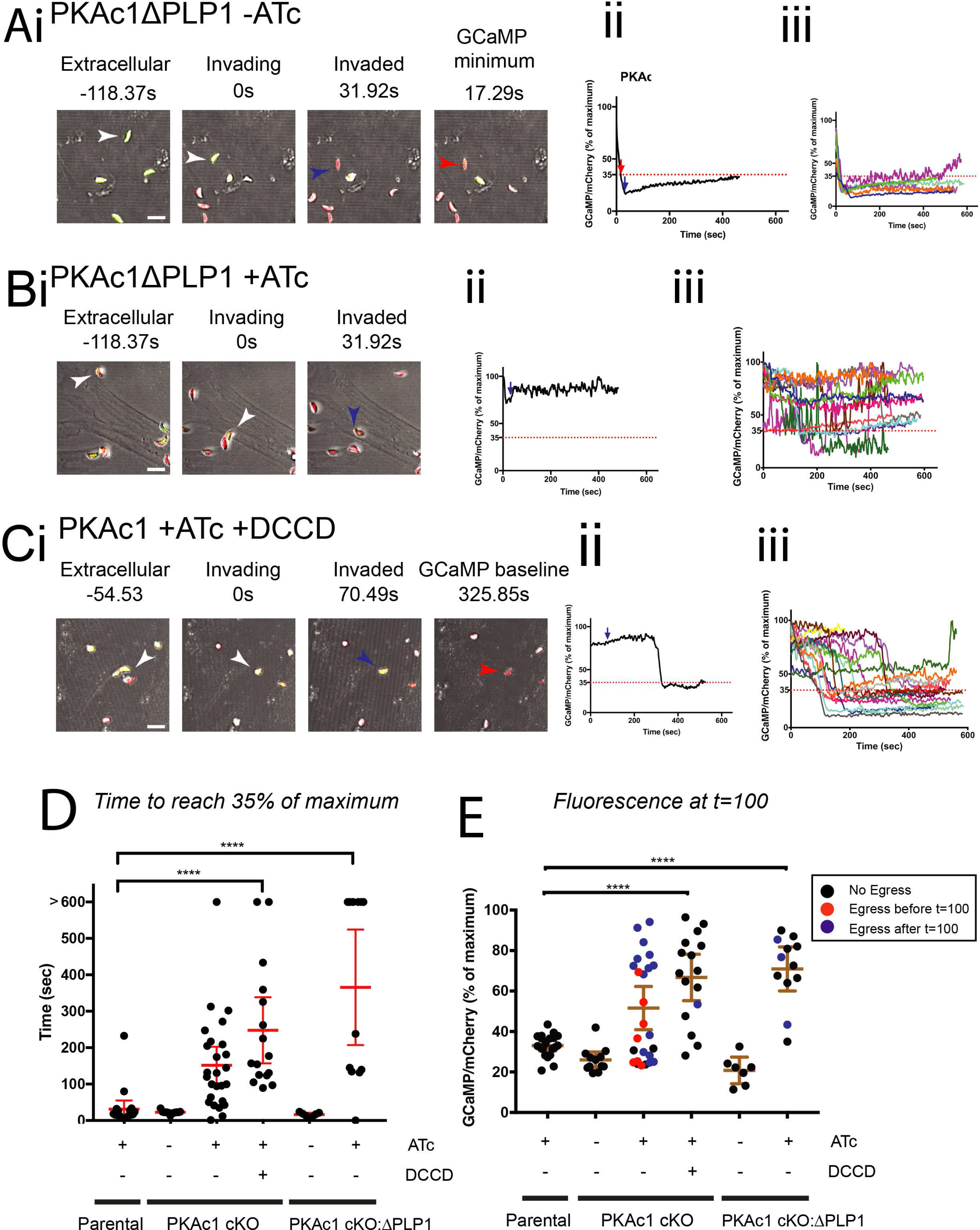
PLP-1 activity does not affect defective Ca^2+^ dynamics upon loss of PKAc1. (A) Live cell imaging of PKAc1 cKO/APLP1 without and (B) with ATc. (C) PKAc1 cKO treated with ATc and 50pM DCCD. All parasite lines are stably expressing GCaMP6f/mCherry at the *uprt* locus. In A, B and C; (i) shows images of individual movies, showing stills when parasites are extracellular, invading, upon completion of invasion (blue arrow) and then when the normalised GCaMP/mCherry ratio, reaches 35% of maximum (see Movies S10, S11 and S12, respectively). (ii) Shows, in each case, a graphical representation of tracked parasites in (i), where blue arrow denotes moment of completion of invasion and green arrow shows moment of host cell egress. Dotted lines marks 35% of maximum GCaMP/mCherry ratio, defined as ‘baseline’. (iii) Shows, in each case, overlays of all traced parasites. Figure S9 shows individual traces of each tracked parasite, for PKAc1 cKO/ΔPLP1 with and without ATc and Figure S10 outlines individual traces of PKAc1 cKO +ATc +50uM DCCD. (D) Graphical representation of each tracked parasite across all conditions showing time to reach 35% of maximum which is defined as ‘baseline’. Parental and PKAc1 cKO lines are from Figure 5, (E) Graphical representation of normalised GCaMP/mCherry ratio at t= 100 seconds across all parasites in all conditions. Data in D and E is represented as mean ± 95% confidence interval. P values are calculated pairwise, using an unpaired two-tailed t-test, where **** =<0.0001.

### PKAc1 negatively regulates the basal cytosolic Ca^2+^ levels

To further dissect the role of PKAc1 in negatively regulating [Ca^2+^]_cyt_ we investigated how the extracellular environment affects cytosolic concentrations of Ca^2+^. First of all, we tested if PKAc1 controls Ca^2+^ transport across the plasmamembrane. To test this, we loaded extracellular tachyzoites with the Ca^2+^-responsive dye Fura-2 and performed calibrated measurements on parasite populations using fluorometry. We found that, when suspended in a Ca^2+^-free buffer, PKAc1-depleted parasites had a statistically higher [Ca^2+^]_cyt_, containing nearly two-fold higher levels (∼33 nM compared with ∼ 58 nM) (Figure 7A). PKAc1 depleted parasites were also observed to have higher [Ca^2+^]_cyt_ than PKAc1 expressing parasites when suspended in salines containing [Ca^2+^] of 1 μM, 100 μM and 1 mM; however, in these cases statistical significance was not reached (Figure 7A).

**Figure 7:**
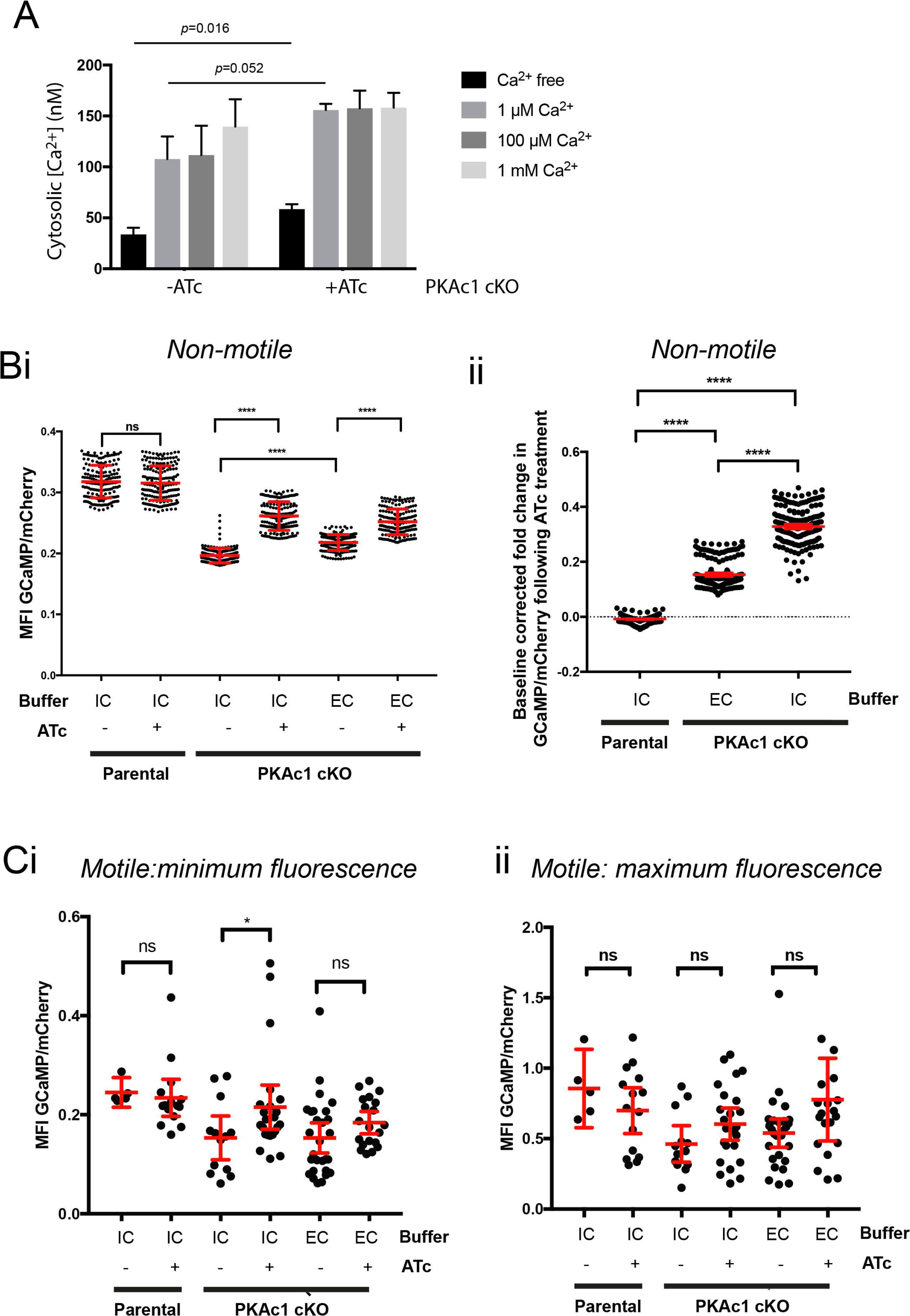
Minimum cytosolic Ca2+ levels are negatively regulated by PKAc1 in extracellular tachyzoites. (A) Measurements of intracellular Ca^2+^ using Fura-2 upon increasing concentration of extracellular Ca^2+^. Data represents mean ±SEM, Statistical significance was assessed using unpaired t-test. (B) Quantitation of GCaMP6/mCherry in Parental and PKAc1 cKO in Intracellular (IC) and Extracellular (EC) buffer with and without ATc treatment in non-motile tachyzoites. (i) represent comparisons of raw ratios beween conditions whereas (ii) represents baseline corrected levels upon treatment with ATc represented as a ratio, where; [Condition +ATc]_intensity_ − [Condition -ATc] _intensity_/[Condition - ATc]_intensity_. Analysis of conditions as in ‘B’ of motile tachyzoites. (i) represents a comparison of minimum GCaMP/mCherry fluorescent levels, whilst (ii) compares maximum levels. All data in B and C is mean ± 95% confidence intervals. P values are calculated pairwise, using an unpaired two-tailed t-test, where ns= not significant, *=<0.05, **** =<0.0001.

Measuring parasites on a population level is unable to distinguish between [Ca^2+^]_cyt_ in tachyzoites that are motile vs non-motile, which are known to differ in Ca^2+^ levels ^9,11^. We therefore used live cell imaging to capture freely moving extracellular GCaMP6/mCherry-expressing tachyzoites and then segregated these based on whether they were motile or not. Given that we observed the biggest effect of PKAc1 in invaded tachyzoites we also wished to dissect the effect of changing [K^+^] concentration, which is considered to be a major environmental change between the intracellular and extracellular environment. To mimic this, we also quantitated motility and corresponding GCaMP6 fluorescence levels when parasites were subjected to extracellular (EC) or Intracellular (IC) buffers which differ in [K^+^].

First, we analysed the [Ca^2+^]_cyt_ in non-motile parasites subjected to IC and EC buffers with and without expression of PKAc1 (Figure 7Bi and ii). Here we found that whilst there was no difference in fluorescence levels of parental parasites upon ATc treatment we found that loss of PKAc1 expression caused a significant increase in [Ca^2+^]_cyt_ in both IC and EC buffers. Indeed, by either comparing the raw GCaMP6/mCherry ratio (Figure 7Bi) or when transforming the data to look specifically at the relative change in fluorescence normalised to before ATc treatment, we found that loss of PKAc1 not only significantly increases resting [Ca^2+^]cyt in non-motile tachyzoites, but this difference increases when parasites are in the high [K^+^] IC buffer (Figure 7Bii). This suggests, that PKAc1 plays an important role in negatively regulating the [Ca^2+^]_cyt_ in non-motile tachyzoites, which is more pronounced when exposed to high extracellular [K^+^], typical of a host cell.

We then analysed [Ca^2+^]cyt in motile tachyzoites, by quantitating minimum and maximum GCaMP6 fluorescence levels and comparing these between parasites in either IC or EC buffer. Quantitating minimum fluorescence levels showed that whilst ATc had no effect on GCaMP6 fluorescence in parental lines, loss of PKAc1 expression led to a statistically significant rise in fluorescence levels only in IC buffer and not EC buffer (Figure 7Ci). Interestingly, we observed no differences in the maximum GCaMP6 fluorescence levels suggesting that PKAc1 does not affect the upper levels of [Ca^2+^]cyt flux in motile tachyzoites (Figure 7Cii). Overall, this analysis strongly suggests that PKAc1 negatively regulates resting [Ca^2+^]_cyt_, levels which is accentuated in IC buffer mimicking the [K^+^] concentration in host cells.

### cAMP levels negatively regulate microneme secretion

Given that PKA is regulated by cAMP we wondered whether levels of this cyclic nucleotide act as a suppressive signal to inhibit motility related processes. To determine if cAMP levels could play a role in negatively regulating egress and motility we applied the membrane permeable analogue 8-Br-cAMP to tachyzoites and quantified the levels of microneme secretion (Figure 8). Cyclic nucleotide analogues are considered not very membrane permeable and thus require large extracellular concentrations to sufficiently leak across membranes. To control for these high concentrations required we used, as a control, 8-Br-cGMP. cGMP has been implicated in the activation of microneme secretion and thus should have the opposite effect of that expected of changes in cAMP levels if these analogues are working specifically.

**Figure 8:**
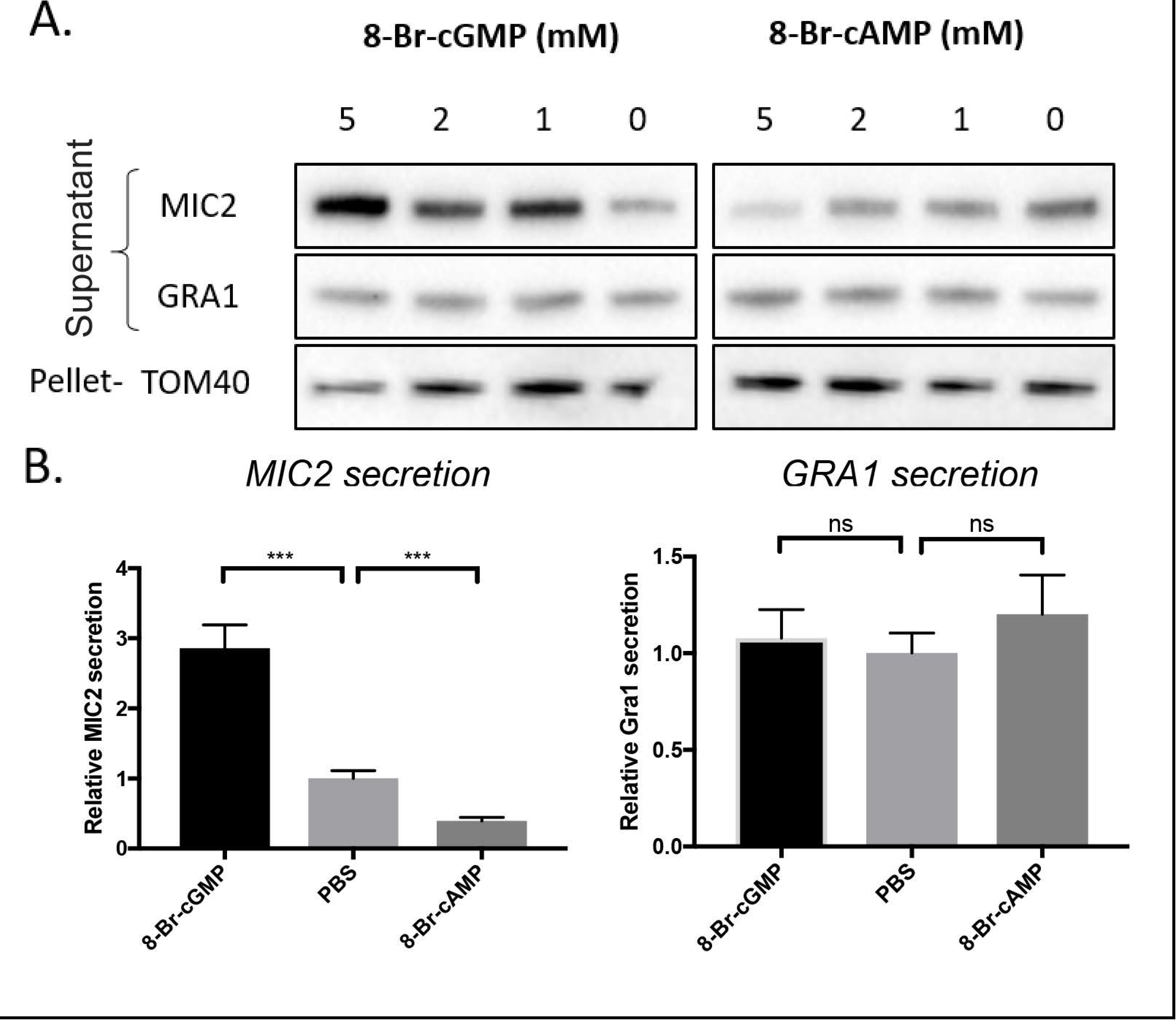
Influence of cAMP and cGMP analogues on microneme secretion in extracellular tachyzoites. (A)Representative secretion assays of wildtype (Δku80:HX) extracellular parasites in response to treatment with 8-Br-cGMP or 8-Br-cAMP (1xPBS as vehicle only control) for 20 minutes. Secretion supernatants were probed for MIC2 and GRA1. TOM40 used as a pellet loading control. (B) Quantitative analysis of secreted proteins over multiple replicates. Parasites treated with either 5mM 8Br-cGMP, 5mM 8Br-cAMP or equivalent 1xPBS solvent control for 20 minutes. Pixel intensity values at 5mM compounds for MIC2 and GRA1 were standardized to TOM40, then normalized to the 1xPBS control which is plotted as 100%, normal constitutive secretion. Y-axis represents relative pixel intensity value, normalized to TOM40 pellet, and again to 1xPBS control. Data represents mean ± SEM. P values are calculated pairwise, using an unpaired two-tailed t-test, where ns= not significant, *=<0.05, **** =<0.0001.

Wildtype extracellular tachyzoites were separated from host cells and treated with increasing concentrations of either 8-Br-cGMP or 8-Br-cAMP. We observed, as expected, that increasing concentrations of 8-Br-cGMP resulted in consistently more secretion of MIC2 into the supernatant (Figure 8A) as previously shown ^40^. Most interestingly, we found that increasing doses of 8-Br-cAMP resulted in a dose-dependent decrease in MIC2 secretion into the supernatant (Figure 8A). Across multiple experiments, we observed a consistent and reproducible effect whereby 8-Br-cGMP results in a relative increase in MIC2 secretion whereas treatment with 8-Br-cAMP results in a relative suppression, whilst levels of the dense granule secretion as measured by Gra1 remained unchanged (Figure 8B, C). This therefore suggests that cAMP has an opposing effect to cGMP in the suppression of microneme secretion.

## Discussion

Critical to the establishment and perpetuation of infection by apicomplexan parasites is their ability to sense and respond to a changing host environment. Environmental cues allow for parasites to undergo differentiation when encountering a new host or to activate motility and egress at an optimal time during host cell infection cycles. Parasites must also switch off motility when invasion of a new host cell is complete in order to start replication. Several environmental factors have been identified that activate differentiation and motility in *Plasmodium* spp. and include mosquito xanthurinic acid ^41^, which activates sexual stage differentiation, low extracellular [K^+^] ^22 42^, which stimulates microneme secretion in blood stages and low pH (high [H^+^]) that activates cell traversal in liver stages ^43^. *Toxoplasma* too activates motility upon a drop in extracellular [K^+^] or pH suggesting that sensing of some environmental cues are conserved across the Apicomplexa ^20,21^. Despite the identification of these environmental cues it is still not understood how apicomplexan parasites transduce these signals across the plasma membrane to induce Ca^2+^-dependent motility and differentiation.

cAMP/PKA signalling in eukaryotes is a major mechanism to relay environmental changes and alter the cellular response. This typically occurs when GPCRs receive extracellular signals and activate adenyl cyclases (AC) to produce cAMP, or alternatively, prevent its breakdown by inhibiting cAMP-specific 3’ 5’-phosphodiesterases (PDEs). In order to see if *Toxoplasma* could also use a similar mechanism to relay environmental cues, we identified a PKA orthologue – PKAc1 – that localises to the parasite periphery. We performed immunoprecipitation of PKAc1 followed by gel- and whole eluate-based identification of interacting proteins and could only robustly identify a likely regulatory subunit PKAr1, suggesting, unlike other eukaryotic systems, *Toxoplasma* PKA does not stably interact with ACs, PDEs or GPCRs. Furthermore, PKAr1 likely derives membrane affinity through myristoylation and palmitoylation at its N-terminus, rather than through interactions with AKAP proteins as in metazoans. Together, this work suggests that PKAc/PKAr either interacts with other proteins in much less stable complexes or that cAMP-dependent signalling in *Toxoplasma* (and other apicomplexan species) is somewhat different from that in metazoans. In this regard, it is also interesting to note that some ACs (as well as guanylate cyclases) in Apicomplexa appear to be directly fused to ion transporter-like domains, suggesting a novel mechanism employed by this group parasites to allow for sensing changes in extracellular ion concentrations and relaying these across the membrane to initiate intracellular signalling cascades ^44,45^.

Controlled depletion of PKAc1 highlighted the importance of this kinase during the *Toxoplasma* lytic cycle and revealed a unique phenotype, whereby tachyzoites rapidly exit the host cell shortly after invasion. In *Plasmodium falciparum*, PKA phosphorylates serine 610 (S610) on the cytoplasmic tail of the adhesin AMA1 and genetic mutation of this phospho-site leads to defects in invasion ^23^. Given these findings in *P. falciparum* and the role that AMA1 plays in formation of the moving junction, we expected to find defects in invasion or sealing of the PVM in *Toxoplasma* PKAc1 mutants. However, using available assays and monitoring the formation of the PVM in real-time using live cell imaging we could detect no apparent defects and instead could show that egress of host cells occurs well after invasion is complete, sometimes on average ∼200 seconds after invasion is complete. Supporting this notion, we demonstrated that loss of PLP-1 reverses loss of PKAc1 during host cell invasion suggesting that aberrant activation of this cytolytic protein in this mutant is responsible for the phenotype. In this regard, it is also interesting to note that the cytoplasmic tail of AMA1 in *Toxoplasma* contains an aspartic acid at residue 557, the residue which is equivalent to S610 in *P. falciparum* ^23^. Aspartic acids share physicochemical properties with phosphorylated serine residues and thus *Toxoplasma* may have lost, or never gained, the capacity to regulate AMA1 tail function by PKA phosphorylation.

Our data suggest that PKAc1, upon invasion, negatively regulates cytosolic Ca^2+^ levels, allowing for the rapid suppression of the Ca^2+^ signal to promote a stable intracellular infection. This suggests that PKAc1-dependent signalling is important for sensing of the host cell environment and suppressing [Ca^2+^]_cyt_ to turn off motility and begin the replicative cycle. [K^+^] is considered to be at a relatively high concentration inside (host) cells and lower in the extracellular environment and previous reports have shown that [K^+^] acts as a major regulator of motility by somehow supressing [Ca^2+^]_cyt_. Using GCaMP6 as a [Ca^2+^]_cyt_ biosensor we were also able to show that higher levels of extracellular [K^+^] resulted in lower levels of [Ca^2+^]_cyt_ as previously reported ^46^. Furthermore, we were able to show that PKAc1 negatively regulates basal [Ca^2+^]_cyt_ levels in both high (IC buffer) and low (EC) [K+]. Most interestingly, however, we found that in high [K^+^] loss of PKAc1 led to a higher level of [Ca^2+^]_cyt_, suggesting that this kinase is especially important for repressing basal levels of [Ca^2+^]_cyt_ when inside the host cell.

We also provide evidence here that cAMP levels negatively regulate microneme secretion. This together suggests that PKAc1 and cAMP signalling, more generally, might act as a signal that parasites use to relay intracellularity and negatively regulate host cell egress. It recently has been shown that PKAc3 in *Toxoplasma* acts as a negative regulator of bradyzoite differentiation ^24^, lending credence to the notion that cAMP signalling may act more generally to keep tachyzoites in the cell cycle and prevent egress from the host cell. For this hypothesis to be true cAMP levels would need to be high in intracellular parasites, diminish nearing host cell egress and remain low during zoite motility. Furthermore, a rapid induction of cAMP would need to occur upon host cell invasion. Given the availability of cAMP biosensors these questions are now imminently addressable and will be interesting to investigate.

Secretion of the perforin-like protein –PLP-1 – from the micronemes and its activation by PV acidification have been shown to both be important for host cell egress ^36 47^ We have demonstrated that early egress by PKAc1-deficient tachyzoites is dependent on PLP-1 and PV acidification but at this stage have not been able to determine whether PKAc1 directly regulates PLP-1 secretion and activity or that they operate in different pathways, both of which are required for egress to take place. Given that PKAc1 negatively regulates [Ca^2+^]_cyt_ in a buffer that mimics the intracellular environment and that artificially increasing cAMP levels can supress microneme secretion, it seems plausible that this kinase may directly regulate PLP1 secretion and activity, but still remains to be directly tested.

Whilst this paper was under review another manuscript was published describing the role of *Toxoplasma* PKAc1 ^48^. Here Jia et al use a conditional dominant negative PKAr, which cannot respond to cAMP and in doing so show that PKAc1 is important for negatively regulating early egress from host cells. Unfortunately, we were unable to robustly look at the role of PKAc1 during egress, due to the time it takes to deplete PKAc1 levels and the stochastic nature of host cell egress. Our work however, is wholly consistent and complementary with their findings and furthermore, also describes the role of PKAc1 in negatively regulating basal [Ca^2+^]_cyt_ levels especially in high [K^+^] like that found in host cells. Indeed, these findings also provide a mechanism as to why dominant negative PKAr from Jia et al can induce early egress and provide evidence as to why there is a preponderance of Ca^2+^ regulated proteins found to be more phosphorylated in the absence of PKAc1 activity ^48^. Jia et al also suggest that PKA and Protein Kinase G (PKG) signalling are connected. Our results our also in line with this as PKG signalling is considered to be connected with Ca^2+^ signalling where we have focussed our study on.

Upon transmission from an infected female anopheles mosquito *Plasmodium* sporozoites must traverse through tissue and set up infection in liver hepatocytes. To achieve this, sporozoites are programmed to traverse through cells, whereby invasion leads to the creation of a transient vacuole and the subsequent PLP-1-dependent egress of the host cell. Work undertaken largely in rodent *Plasmodium* species suggests that the decision to traverse a cell is made upon interaction with the CD81 surface receptor, rather than occurring after sporozoite invasion is complete ^43,49^. *Toxoplasma* parasites described here lacking PKAc1 behave in an analogous manner to a traversing sporozoite, whereby a PVM is formed and then egress is induced in a PLP-1-dependent manner. Our work therefore opens up the possibility that *Plasmodium* sporozoites may also be able to regulate productive invasion or cell traversal using cAMP signal transduction pathways. Clearly, this will be interesting to investigate in the future.

## Materials and Methods

### Plasmid construction

Detailed methodologies and oligonucleotides used for construction of plasmids are available in Supplementary Information.

### Toxoplasma transfection and in vitro culture

*Toxoplasma gondii* tachyzoites were cultured under standard conditions. Briefly, Human Foreskin Fibroblasts (HFF)(ATCC), Vero and immortalised MEFs expressing membrane-bound tdtomato ^35^(a kind gift from C. Allison, WEHI) were grown in DME supplemented with 10% heat inactivated Cosmic Calf Serum (Hyclone). When infecting HFFs with tachyzoites media was exchanged to DME with 1% FCS. All cells were grown in humidified incubators at 37°C/10% CO_2_. Transfection proceeded using either a Gene Pulser II (BioRad) or a Nucleofector 4D system (Lonza). Gene Pulser II transfection took place using standard procedures using 50 µg of purified DNA and the DNA was linearized if seeking homologous integration ^50,51^. Nucleofection proceeded by using 2 × 10^6^ tachyzoites and 5 μg of DNA in 20 μl of buffer P3 (Lonza) and pulsed using program F1-115.

### Plaque assays

Plaque assays were performed by inoculating 100-500 tachyzoites onto confluent monolayers and leaving the cells undisturbed for 7-8 days. Monolayers were then fixed in 100% ethanol and stained with crystal violet (Sigma).

### Immunofluorescence Assay (IFA)

IFA’s of *Toxoplasma-infected* host cells were undertaken using standard procedures detailed in Supplementary Information. Images were captured on an AP DeltaVision Elite microscope (GE Healthcare) equipped with a CoolSnap2 CCD detector and captured with SoftWorx software (GE Healthcare). Images were assembled using Image J and Adobe Illustrator software. Ty (mAb BB2), HA antibodies (3F10)(Roche), GAP45 ^52^ and ISP1 ^53^ were all used at 1:1000 in 3% BSA/PBS. Secondary Alexa Fluor conjugated antibodies (Molecular Probes) were all used 1:1000.

### Host cell Invasion and attachment assays

Detailed protocols used for monitoring both Invasion and host cell attachment are available in the supplementary text. The invasion assay proceeded, largely as previously described ^54^. Briefly, Parasites lines were treated ±ATc and resuspended in intracellular buffer (IC) to prevent invasion and left to spun down onto the HFF monolayer in an 8-well chamber slide (Ibidi). IC buffer was exchanged for DME/1%FCS/10mM HEPES pH 7.5 and incubated for 10 minutes at 37°C. Samples were then, fixed, stained with anti-SAG1 mAb 6D10 (1:2000), permeabilized and stained with rabbit anti-GAP45 (1:1000)^52^ followed by secondary antibodies then imaging on a Ziess Live Cell Observer and quantitation by manual counting. Invasion assays with DCCD were carried out as described above except that parasites were allowed to invade in D1/Hepes medium containing 50 μM DCCD (Sigma).

### Host cell Attachment assays

Detailed protocol for attachment assays is described in the supplementary text. Tachyzoites were resuspended in DME/1% FCS/10mM HEPES pH7.5 at a concentration of 1x 10^7^ tachyzoites/ml. 200μl of tachyzoites was added to each well of a 8-well chamber slide (Ibidi) that contains HFFs fixed with 2.5% formaldehyde/0.01% glutaraldehyde. Parasites were allowed to attach for 30 minutes at 37°C, before fixing again and a standard IFA performed as above using SAG1 or GAP45 antibodies.

### Immunoprecipitation and proteomics

Egressed parasites from three T150 flasks for each of Δ*ku80*:TATi (parental) and Δ*ku80*:TATi:*PKAc1* cKO parasites were resuspended in 1 ml lysis buffer (100 mM HEPES, pH 7.5; 1% NP-40; 300 mM NaCl; 1 mM MgCl_2_; 25 U/ml Benzonase [Novagen] and protease inhibitors without EDTA [Roche]) and incubated on ice for 30 min, before centrifuging for 15 min at 14 000 rpm and 4°C to pellet cellular debris. The lysates were added to 100 μl of anti-HA agarose beads (50% slurry [Sigma]) that had been washed twice with 1 ml PBS/0.5% NP- 40) and incubated for two and a half hours at 4°C, with rotation. The beads were then pelleted by centrifugation at 10,000 x *g* for 15 s and the supernatants removed. The beads were washed 5 times with ice-cold 1 ml PBS/0.5% NP-40. Analysis was performed by running eluates on a NuPAGE 4-12% non-reducing Bis-Tris Gel (Invitrogen) and stained with SyproRuby according to the manufacturer’s instructions (Invitrogen), followed by Page Blue staining (Fermentas). The two prominent bands from the Δ*ku80*:TATi:PKAc1 cKO IP were cut out and identified by mass spectrometry. Whole eluates were also subjected straight to mass spectrometry without protein gel separation and quantitated using a custom label-free pipeline. For more details on procedures and SILAC labelling used for quantifying protein release from micronemes please refer to Supplementary Information.

### Live Cell Imaging

8-well cell culture-treated chambers slides (Ibidi) were seeded with either HFF or ROSA MEFs and grown as outlined above. *Toxoplasma* tachyzoites were treated overnight with 1μg/ml of ATc and harvested as stated above. Tachyzoites were then resuspended in Endo buffer and spun down onto the host cells and then taken to the imaging station.

Tachyzoites and host cells were then imaged using a Leica SP8 confocal, equipped with resonant scanner. For quantitative fluorescent measurements GCaMP was excited with 488 nm krypton/argon laser line, whilst mCherry and tdtomato were excited using the 594 nm and 561 nm laser lines, respectively. All imaging took place with pinhole at 1 Airy unit, typically over 5 z-stacks covering 5 µm. Image analysis took place using ImageJ using in-built and custom macros. Data were processed in Excel and graphed in Prism.

### Intracellular Ca^2+^ measurements

PKAc1 cKO parasites grown in confluent HFFs for 24 h were treated with 1 μg/mL ATc (or the equivalent volume of 100% EtOH; solvent control) for 6 h. Following this, parasites were harvested by passage of cultures through a 26-gauge needle and centrifuged (1500 × *g* for 10 min at 4°C). The parasites were then resuspended in fresh culture medium containing either ATc (final concentration 1 μg/mL) or the equivalent volume of 100% ethanol and added to flasks containing confluent HFFs for up to 24 h. The parasites were then harvested from their host cells and loaded with Fura-2 as described by McCoy et al (2017). Cytosolic Ca^2+^ measurements were performed at 37°C using a PerkinElmer LS 50B Fluorescence Spectrometer as described previously ^55^.

### Microneme secretion assays

Secretion assays were performed and quantitated as described previously ^40^. 6-Br-cAMP and 6-Br-cGMP analogues were solubilized in 1xPBS, and an equal volume of 1xPBS was used as a vehicle only control. Cells were then shifted to 37°C and allowed to secrete for 20, or 60 min as indicated. Secretion was arrested by placing cells on ice for 2 min. Parasites were separated from the soluble secreted proteins by centrifugation at 8000 rpm, 4°C for 2 min. 85μL of supernatant was removed and centrifuged again to remove any remaining cells, and 75 μL of supernatant was removed and boiled with reducing Sample Buffer. Parasites pellet was washed with PBS and boiled in reducing Sample Buffer. Secreted proteins were analysed using antibodies to indicated factors by western blot: TOM40 (1:1000) ^56^, GRA1 (1:2000)^57^, MIC2 (1:5000)^58^, AMA1 CL22 (1:1000) ^59^, SUB1(1:1000) ^60^ and PLP1 ^37^.

## Acknowledgements

We are most appreciative of ongoing help from Kelly Rogers and Lachlan Whitehead (WEHI) for helping with image acquisition and analysis. We would like to thank Giel van Dooren (Australian National University) and Dominique Soldati-Favre (The University of Geneva) for plasmids and antibodies. We would also like to acknowledge Vern Carruthers (University of Michigan) and David Sibley (Washington University) for providing us with antibodies. We are also indebted to Cody Allison (WEHI) for providing us with immortalised tdtomato:ROSA26 MEFs. This work was supported by NHMRC project grants APP1025598 (CJT), APP1047806 (CJT) and an ARC Future Fellowship FT120100164 awarded to CJT CYB and NJK are supported by Agence Nationale de la Recherche (LABEX PARAFRAP ANR-11-LABX-0024) and CNRS-INSERM-Fondation FINOVI (Atip-Avenir Program Project Apicolipid).. We are also grateful for institutional support from the Victorian State Government Operational Infrastructure Support and the Australian Government NHMRC IRIISS. AML is the recipient of an Australian Research Council Discovery Early Career Researcher Award (DE160101035)., AFW is supported by MRC (MR/M011690/1). L.Y is a recipient of a China Council Scholarship.

## Author contributions

A.D.U, M.W, R.J.S, L.F.D, L.Y, E.A.M, N.J.K, S.V.H, M.J.C, A.M.L, R.F.W, A.I.W and C.J.T designed experiments and analysed data. A.D.U, R.J.S and C.J.T managed the project and C.J.T wrote the manuscript. A.D.U, R.J.S, L.Y generated transgenic lines and phenotyped resulting mutants. E.A.M, M.W, C.J.T and R.J.S performed live cell imaging and phenotyped mutant lines. N.J.K and R.F.W performed quantitative microneme secretion assays. S.V.H and A.M.L performed Fura-2 measurements of cytosolic Ca^2+^ levels. L.D and A.I.W performed quantitative mass spectrometry.

---

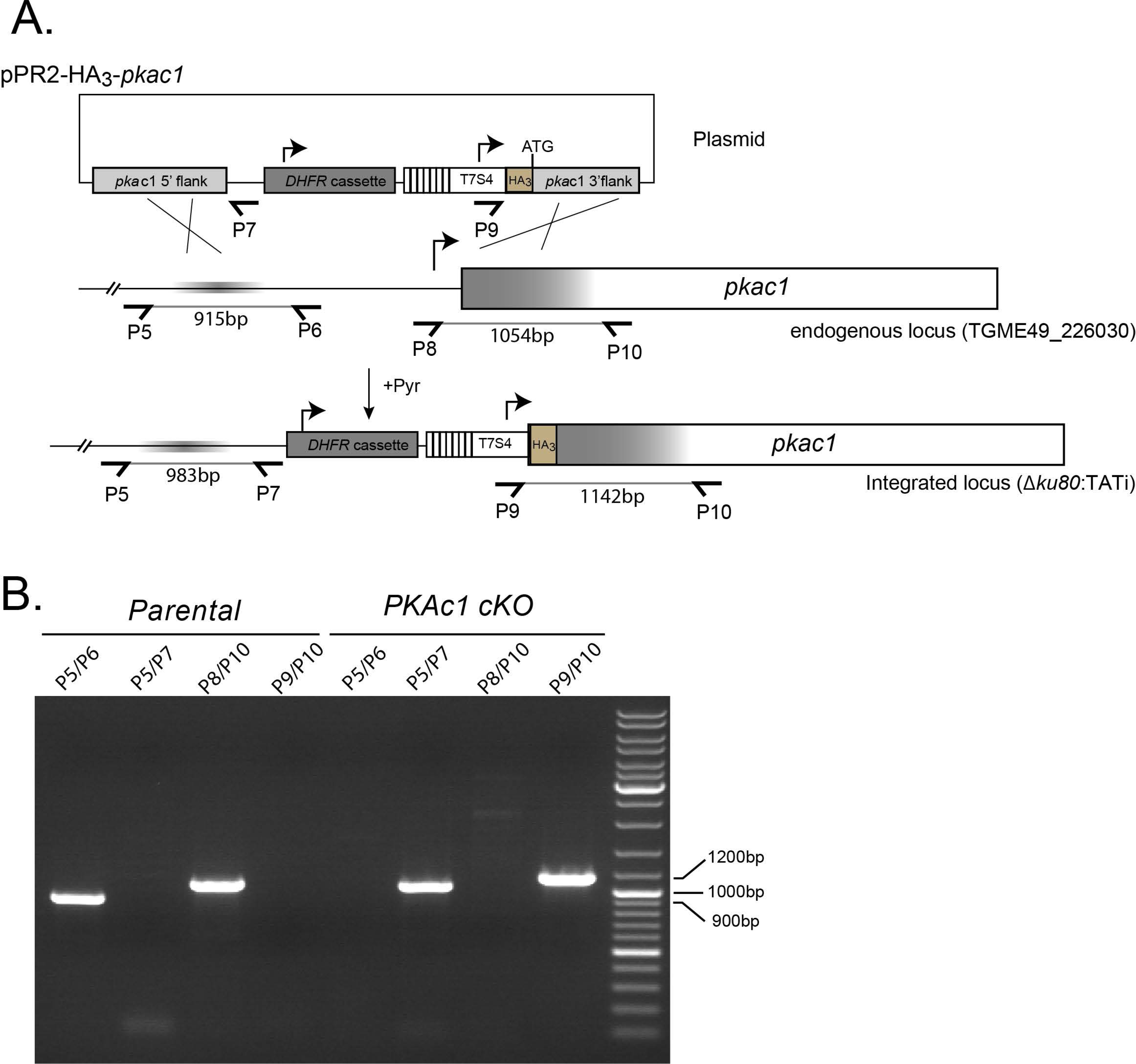

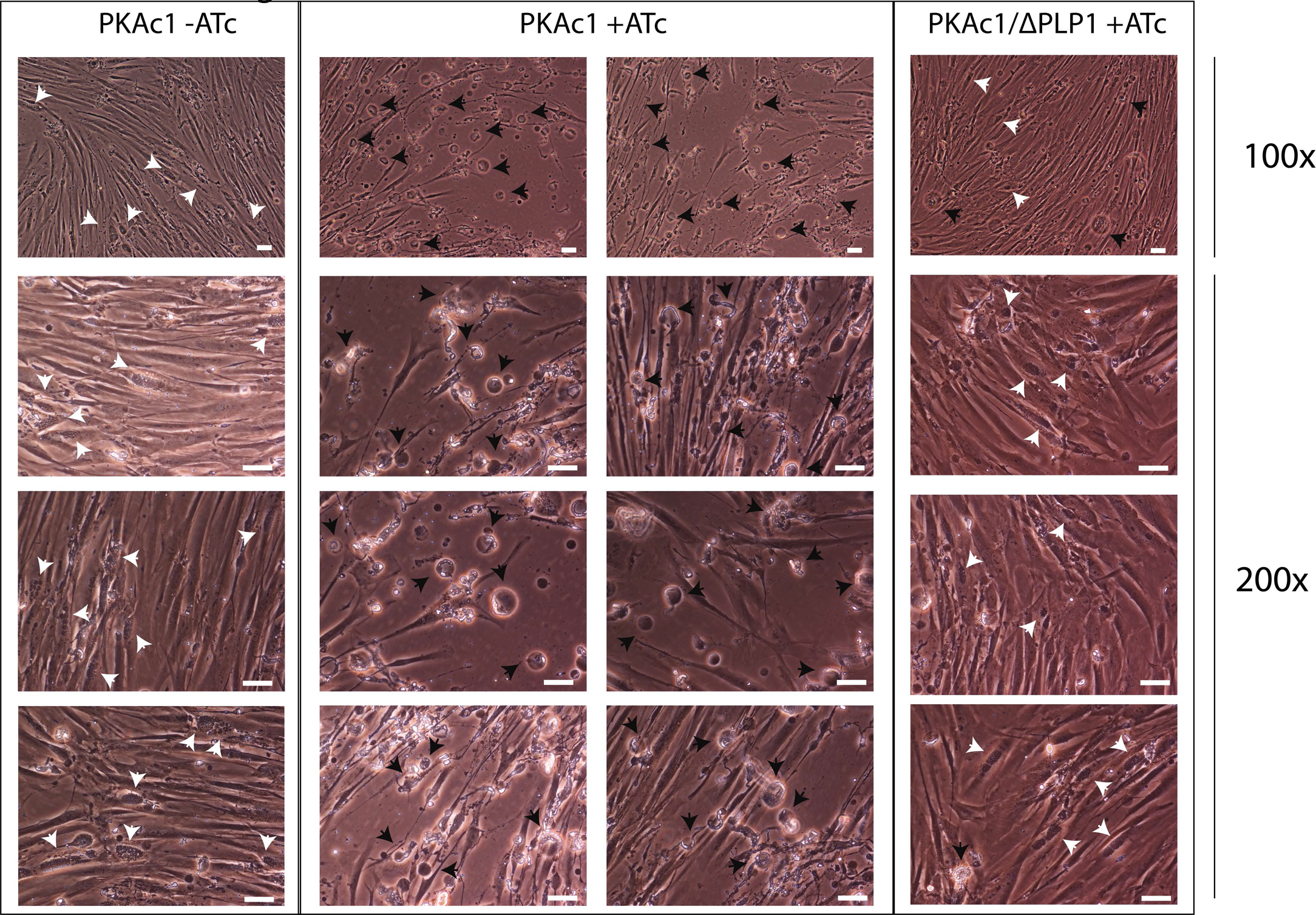

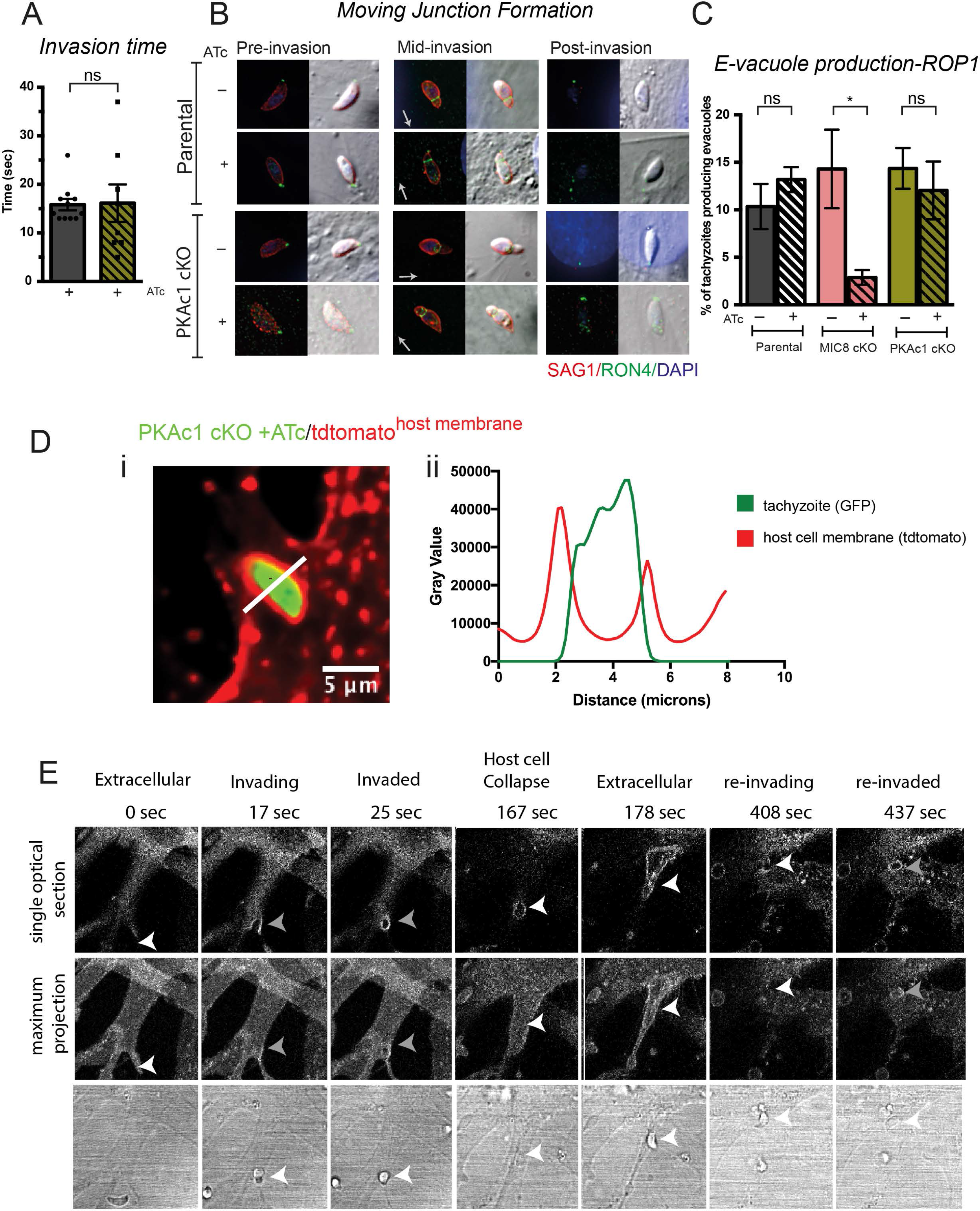

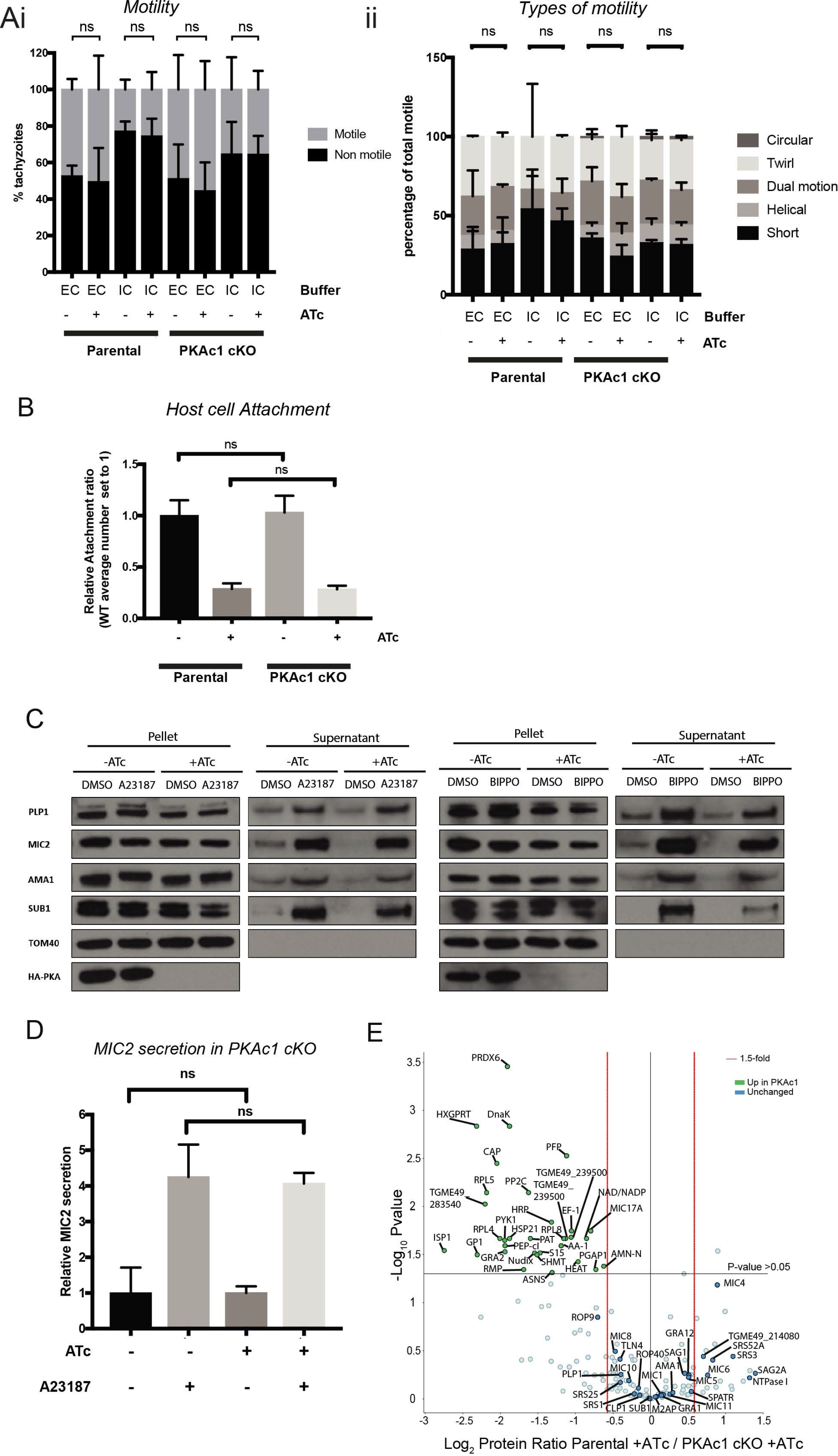

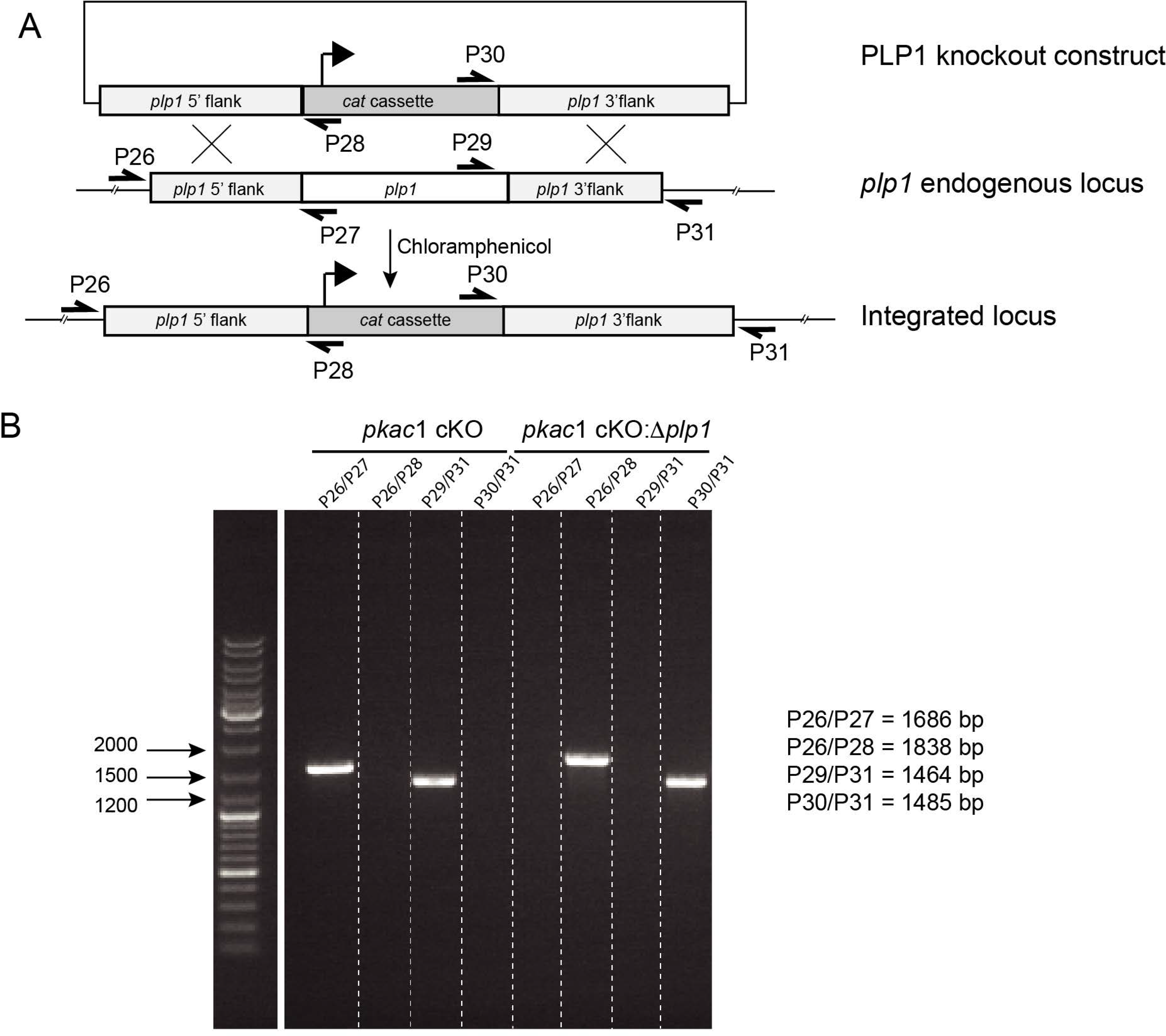

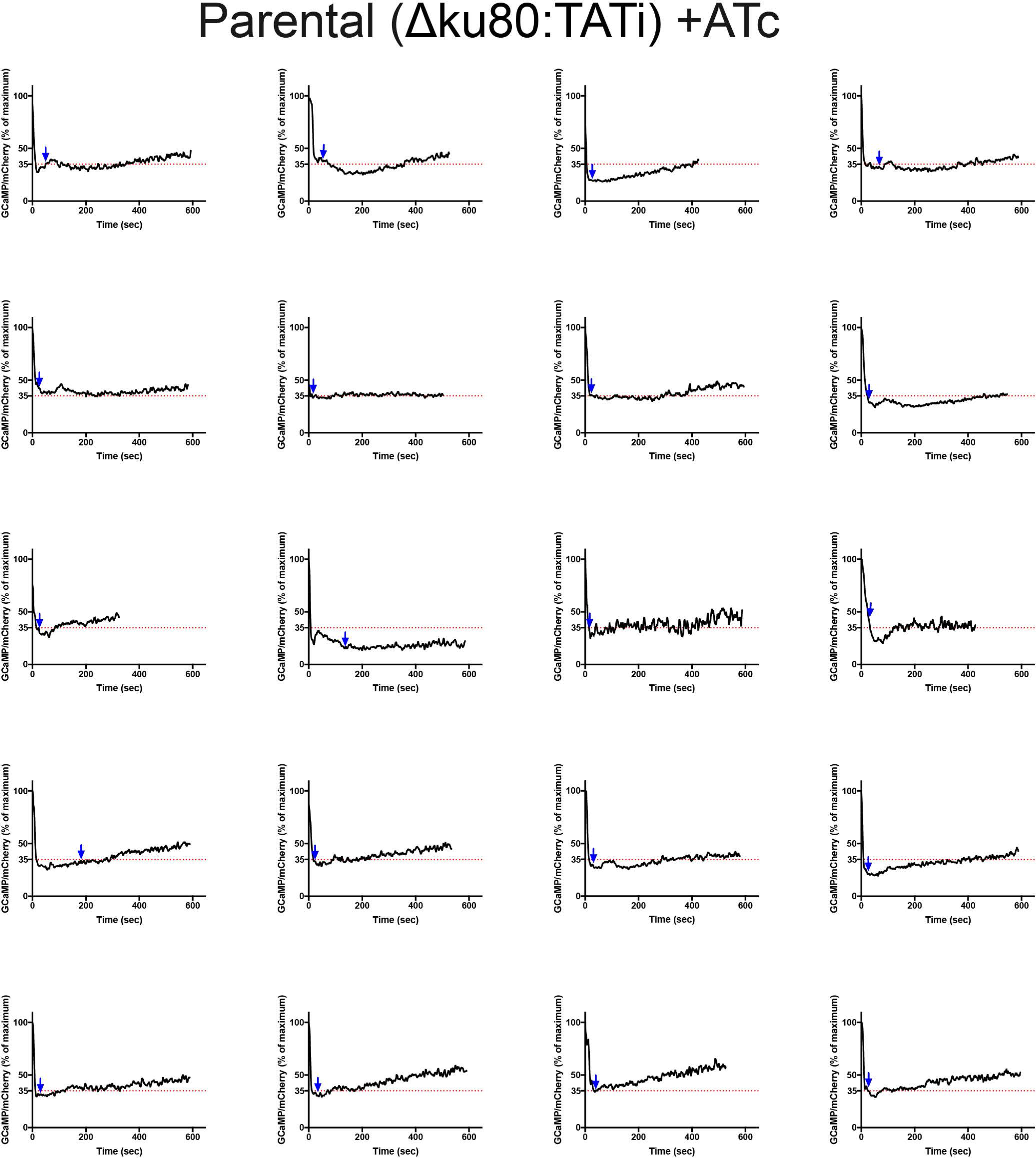

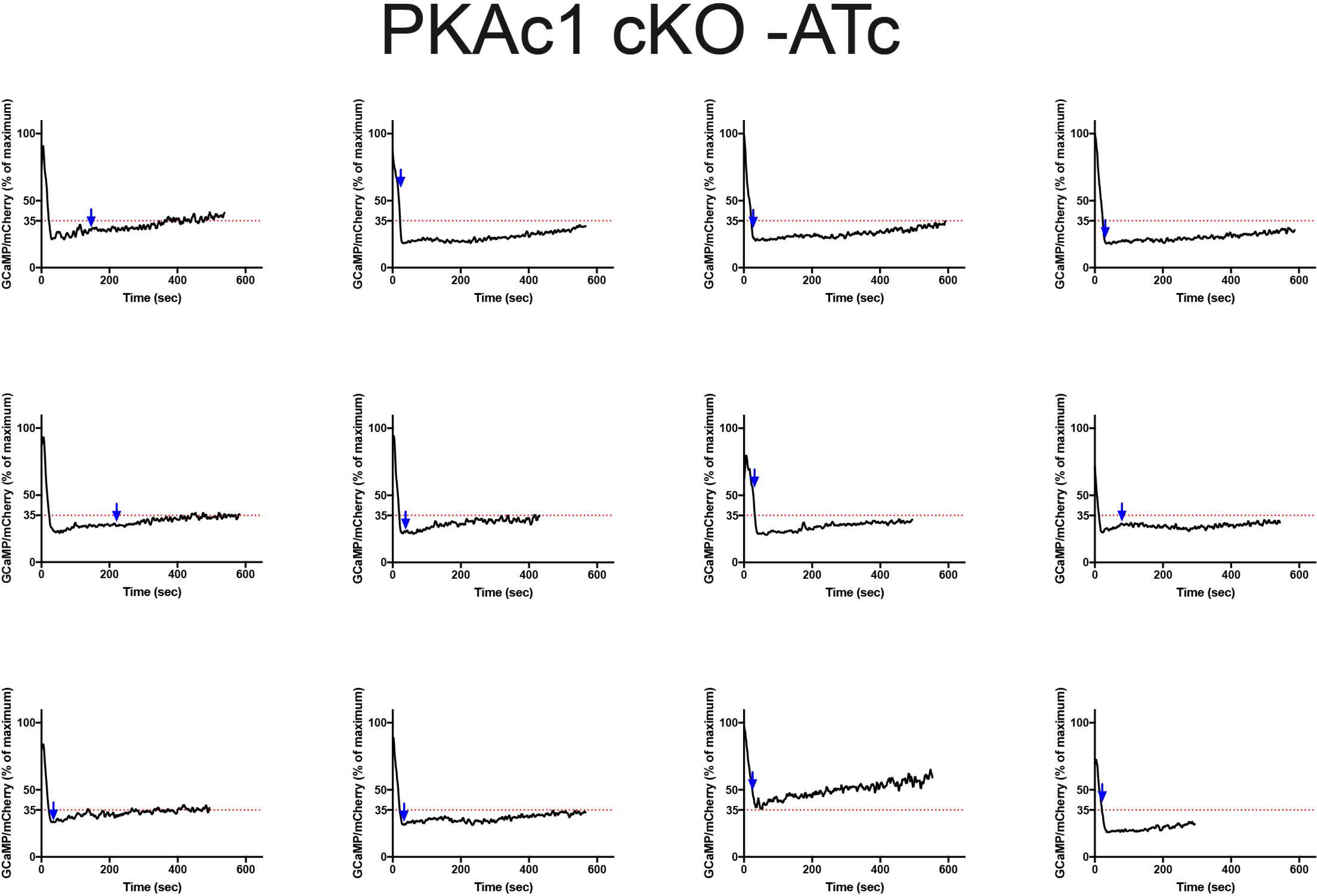

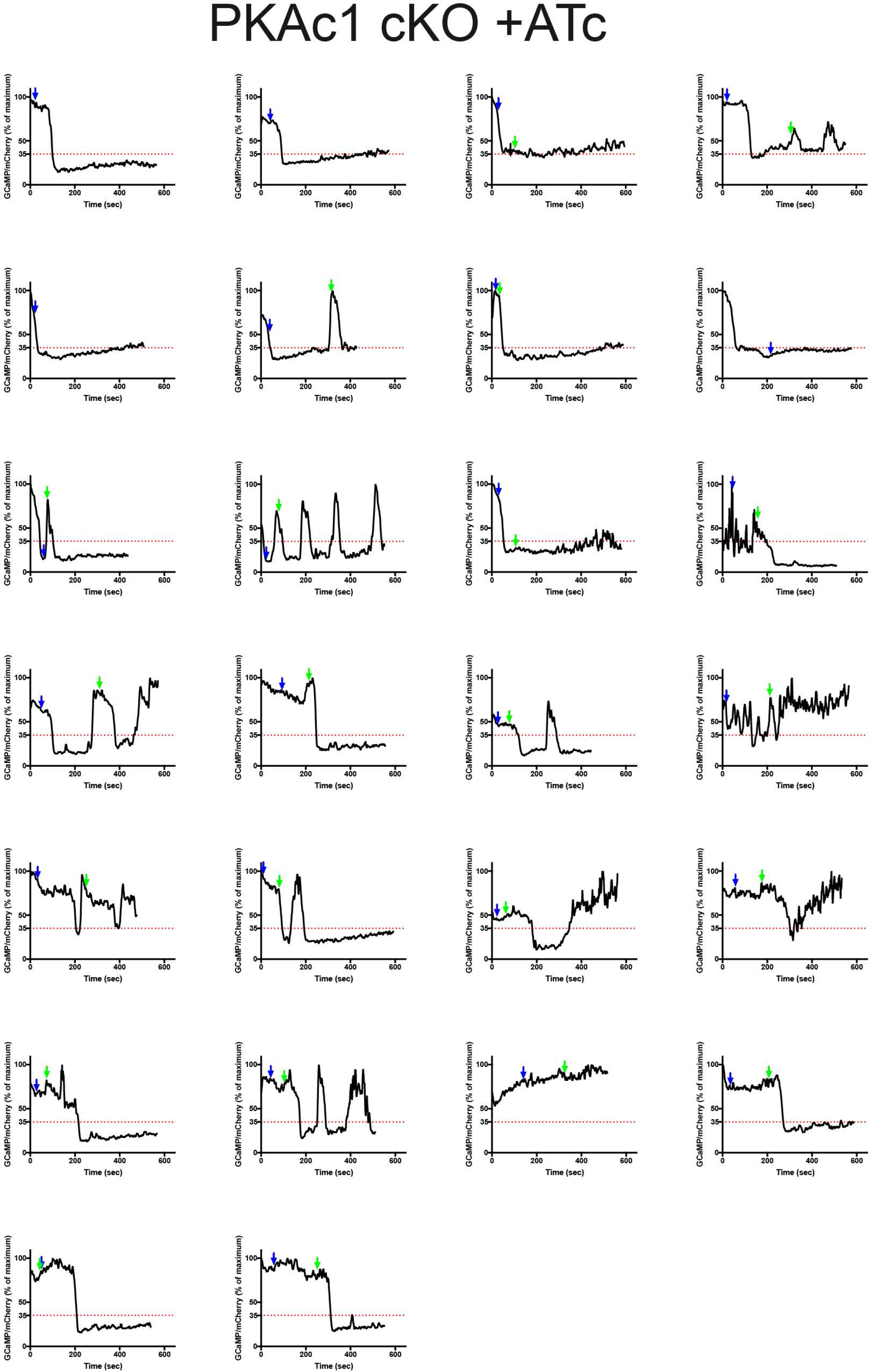

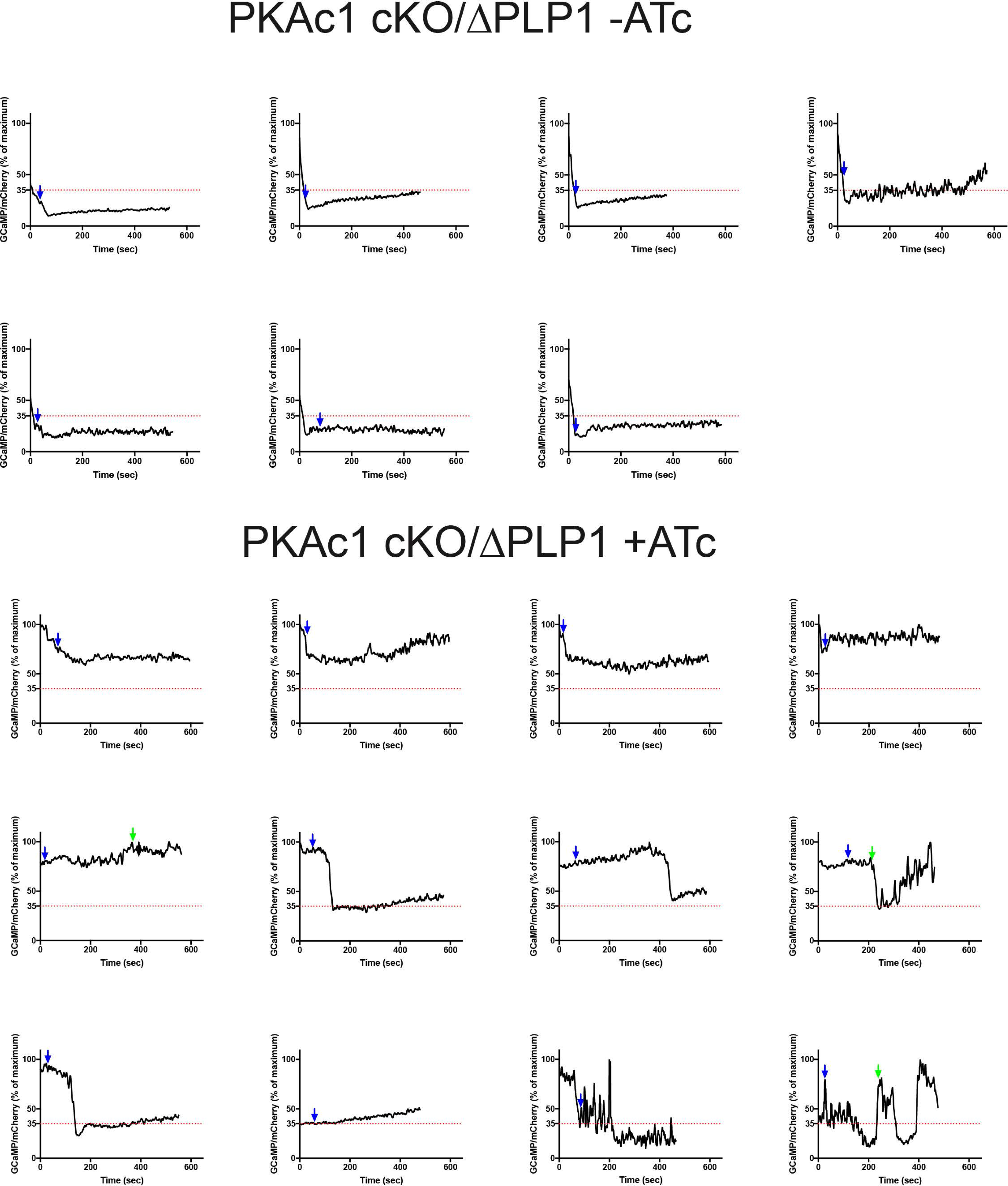

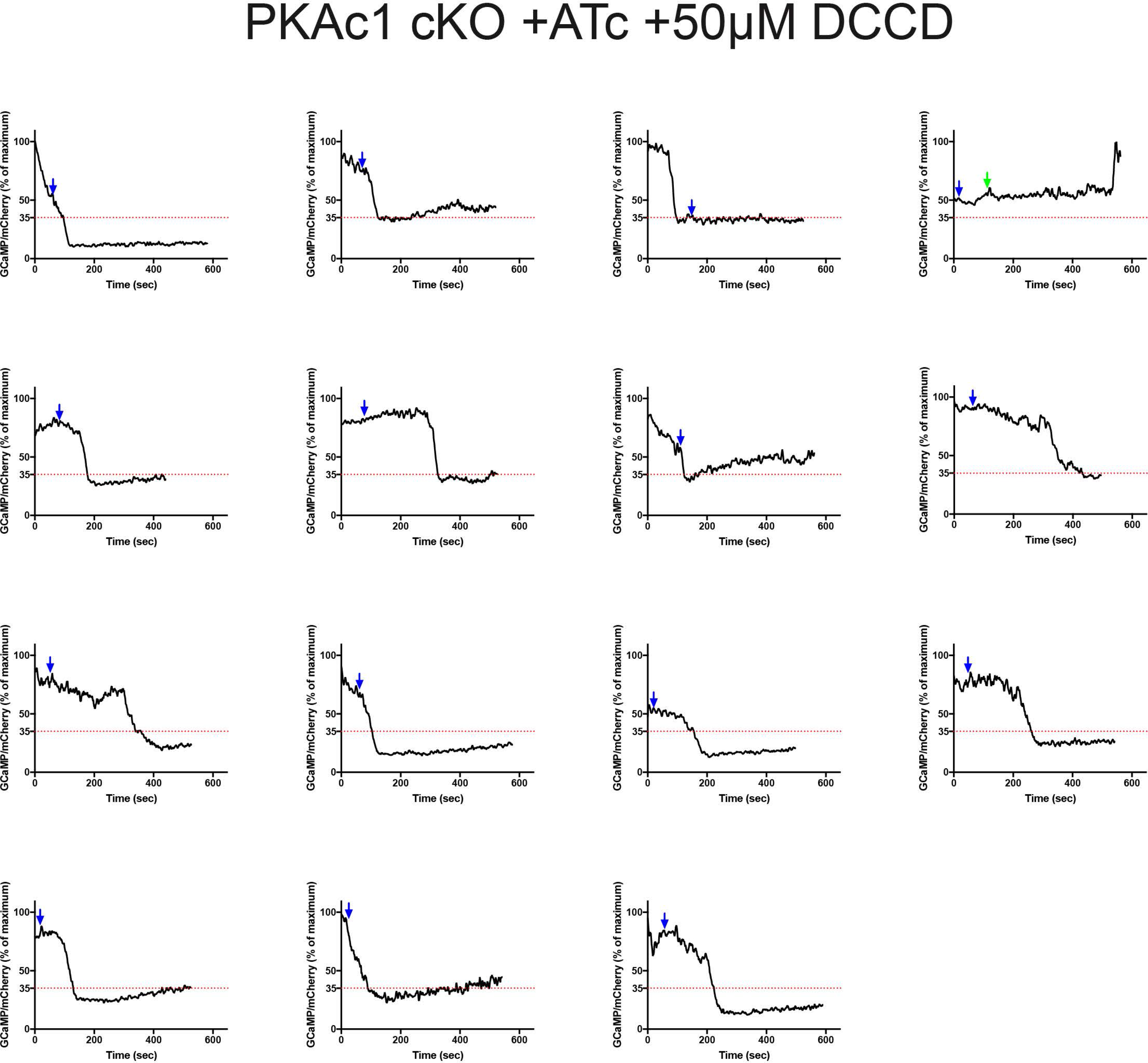

